# Decoding human *in vitro* terminal erythropoiesis originating from umbilical cord blood mononuclear cells and pluripotent stem cells

**DOI:** 10.1101/2023.04.20.537628

**Authors:** Xiaoling Wang, Wei Zhang, Siqi Zhao, Hao Yan, Zijuan Xin, Tiantian Cui, Ruge Zang, Lingping Zhao, Haiyang Wang, Junnian Zhou, Xuan Li, Wen Yue, Jiafei Xi, Zhaojun Zhang, Xiangdong Fang, Xuetao Pei

**Affiliations:** Stem Cell and Regenerative Medicine Lab, Beijing Institute of Radiation Medicine, Beijing,100850, P. R. China; Beijing Institute of Genomics & China National Center for Bioinformation; Institute for Stem Cell and Regeneration, Chinese Academy of Sciences, Beijing 100101, P. R. China; South China Research Center for Stem Cell & Regenerative Medicine South China Institute of Biomedicine, Guangzhou, 510005, P. R. China; College of Life Sciences; Savaid Medical School; Sino-Danish College; School of Future Technology, University of Chinese Academy of Sciences, Beijing 100049, P. R. China; Beijing Key Laboratory of Genome and Precision Medicine Technologies, Beijing 100101, P. R. China

**Author notes:** Correspondence (X.P.); (X.F.); (Z. Zhang); (J.X.). X.W. and W.Z. contributed equally in this work.

**Keywords:** sc-RNA seq, umbilical cord blood mononuclear cells, pluripotent stem cells, *in vitro* erythropoiesis

## Abstract

Ex vivo RBC production generates unsatisfactory expansion, β-globin expression, and maturation of erythroid cells. The underlying mechanisms behind these limitations and ex vivo terminal erythropoiesis from different origins are largely unexplained. In this study, we mapped an atlas of ex vivo terminally differentiated cells from umbilical cord blood mononuclear cells (UCBMNs) and pluripotent stem cells (PSCs), and observed the differential regulatory dynamics of erythropoiesis from these two origins at a single-cell resolution. We detected the presence of hematopoietic stem progenitor cells (HSPCs), erythroid progenitor (e.g., CFU-E), and non-erythroid cells (e.g., macrophages) in the terminal populations. We observed that UCBMN-derived erythropoiesis is more active than PSC-derived erythropoiesis in terms of the cell cycle, stress erythropoiesis, and autophagy at single cell resolution, which may provide new insights into the limitations in cell expansion, globin expression, and maturation in ex vivo RBC production, respectively. We verified that a stress-erythropoiesis-related gene, *TRIB3*, increases the expression of globin genes in ex vivo erythropoiesis. As the major unexpected component detected in terminally differentiated cells, CFU-E were further characterized as having high- or low- expansion capacity based on CD99 expression, which generally decreased over erythropoiesis. By inhibiting CD99 gene expression using antagonists, we increased reticulocyte production in the population. Heterogeneous CFU-Es also exist in bone marrow. Moreover, decreased CD99 expression mediates the interactions between macrophages and CFU-E during ex vivo erythropoiesis. Overall, our results provide a reference for facilitating the development of strategies to improve ex vivo RBC regeneration.

**Highlights:**

- We performed scRNA-seq and cell typing of late stage UCBMN- and PSC-derived cells
- Stress erythropoiesis, autophagy and cell cycle related gene expression different in two origins
- CD99^high^ progenitor cells are a proliferating colony forming unit erythroid subpopulation

## INTRODUCTION

Immortalized cell lines, such as pluripotent stem cells (PSCs), are ideal, stable, and are theoretically infinite resources for red blood cell (RBC) generation *in vitro*. However, PSC-regenerated erythrocytes often have extremely low enucleation ratios and insufficient β-globin expression (Olivier et al., 2016; Wang et al., 2017). Since they possess multi-lineage potency, umbilical cord blood mononuclear cells (UCBMNs) have become one of the most valuable primary cell sources for RBC regeneration. Compared with PSCs, UCBMN-derived erythroid progenitors possess higher proliferation capacities, and the derived RBCs show higher enucleation ratios and β-globin expression levels (Rallapalli et al., 2019). Recently, comparative bulk RNA-seq analysis between embryonic stem cell (ES)- and UCB-CD34^+^-derived erythroid cells provided insights into the limited expansion and defective enucleation of ES-origin RBCs(Wang et al., 2022), but lacked mechanical investigations of ex vivo erythropoiesis at higher resolutions. The mechanisms underlying ex vivo erythropoiesis limitations remain unclear.

Single-cell sequencing technology analysis can considerably improve our understanding of erythropoiesis and the contribution of each cell type within cell populations (Jacobsen and Nerlov, 2019). A previous study constructed a hematopoietic development model characterized by cell population heterogeneity at early stages of erythropoiesis using single cell sequencing technology (Buenrostro et al., 2018). Xin et al. determined specific regulatory networks that controlled ex vivo iPSC-derived erythropoiesis in embryoid body (EB) culture (Xin et al., 2021), providing critical clues to improve iPSC-derived ex vivo erythropoiesis, but this study focused on very early developmental erythropoietic stages. A previous study assessed the transcription dynamics of terminally differentiated human erythroblasts from human CB and bone marrow cells at single cell resolutions (Huang et al., 2020), especially focusing on the mechanisms of in vivo terminal erythropoiesis in ortho-E from two primary sources. Currently, single-cell transcriptomic differences between UCBMN-derived erythropoiesis (UCB-E) and PSC-derived erythropoiesis (PSC-E) at terminal stages and their underlying regulatory mechanism remain unknown.

In this study, we comprehensively mapped and compared the atlases of ex vivo UCBMN- and PSC-derived terminally differentiated cells, and explored the mechanisms underlying limitations associated with ex vivo RBC production (erythroid progenitor expansion, globin expression, and maturation). We characterized the major components, especially CFU-E and macrophages, in the cell populations, and provided various potential strategies for facilitating ex vivo erythropoiesis. We aimed to clarify mechanisms in vitro terminal erythropoiesis and ultimately, to improve existing RBC generation strategies.

## RESULTS

### Cell composition of terminally differentiated erythroid cells derived from UCBMNs and PSCs

We applied the “spin embryonic body” and three-stage differentiation methods to generate mature erythroid cells *in vitro* (Figure S1A, S1B). We comparatively analyzed UCB-Es and PSC-Es at the single-cell transcriptome level on days 21 and 23 of the experiment, when they all reached the highest of CD71 and CD235a double positive, to evaluate differences in their capacity to generate RBCs and the underlying mechanisms (Figures 1A-B). We observed that PSC-E accumulation was lesser than that of UCB-E, which is consistent with previous studies (Galat et al., 2018; Tao et al., 2017). Moreover, PSC-E cells lacked beta/gamma-globin, were relatively larger in size, and showed impaired enucleation when compared to UCB-E (Figures 1C-D).

**Figure 1.**
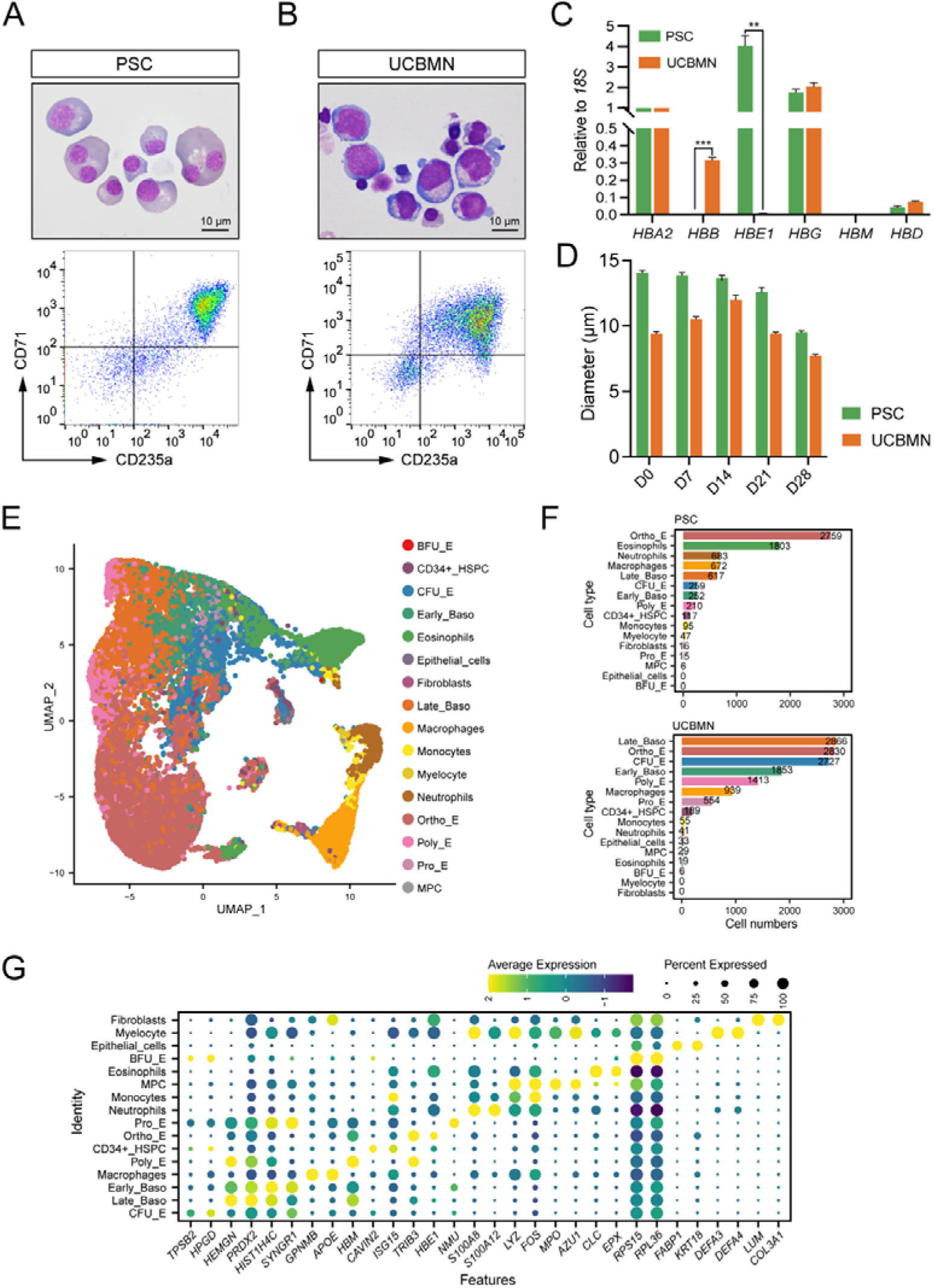
Atlas of PSC- and UCBMN-derived terminal differentiated cells. (A-B) Characterization of sequencing samples by Giemsa staining (upper panels) and flow cytometry analysis on CD71/CD235a (lower panels). Two origin generated erythroid cells reach more than 90% CD71/CD235a double positive in the sequencing time, the cultured day 21- and day 23- erythroid cells derived from UCBMN and PSC cells. (C) Hemoglobin expression relative to *18S* in UCB-E (Day 21) and PSC-E (Day 23) determined by RT q-PCR. The expression of HBB and HBE between two origins shown significant difference (n=3). **: P < 0.01, ***: P < 0.001. (D) Diameters statistical plot of PSC- and UCBMN-derived cultured erythroblasts at different timepoints. Generated cells from two resources all reached about 10µm while PSC-derived cells always much smaller than UCB-derived ones (n=3). (E) The single cell UMAP plot of UCB- and PSC-derived samples in D21 (UCB) and D23 (PSC). 16 cell types are indicated by colors. Based on published data [14, 15] and the Human Primary Cells Atlas (HPCA) database, celltypes were annotated by R package SingleR. (F) Bar plots showing numbers and distributions of each cell type in samples derived from PSC (upper) and UCBMN(lower), respectively. (G) Dot plot is showing the expression levels of top 2 signature genes (ordered by averaged log2 fold change, P value < 0.05) in each cell type. Colors are indicating the scaled average expression of signature genes. Dot sizes are indicating the percentage of cells which are expressing the signature genes in each cell type cluster.

After quality control, 21,105 high-quality single cells were generated for downstream analysis. The datasets we obtained were comparable (Figures S1D-S1H). Analysis of published data (Hu et al., 2013; Li et al., 2014) and the Human Primary Cells Atlas (HPCA) database based on SingleR (Aran et al., 2019) generated 16 reliable cell populations (Figures 1E-G, S1I-S1K). The cells were classified as progenitors/precursors [e.g., CD34^+^ HSPC, BFU-E, CFU-E, and myeloid progenitor cells (MPC)], erythroid cells [e.g., pro-erythroblast (Pro-E), early and late basophils (Baso-E), polychromatic erythroblasts (Poly-E), orthochromatic erythroblasts (Ortho-E)], or non-erythroid cells (e.g., eosinophils, macrophages, neutrophils, monocytes, myelocytes, epithelial cells, and fibroblasts). Erythroid cells dominate the UCB-Es as expected, while erythroid and non-erythroid cells dominate the PSC-Es. Interestingly, CFU-Es and macrophage were present in both systems with relatively higher percentages, while more non-erythroid hematopoietic cells (e.g., eosinophils, neutrophiles) were present in the PSC-E but not the UCB-E (Figures 1F, 1G). Taken together, sc-RNA seq reveals the diversity of terminally differentiated cells from both PSC- and UCBMN-derived sources.

### UCBMN- and PSC-derived terminal erythroid differentiation are differentially and dynamically regulated

The cell compositions of terminally differentiated cells indicated that PSC- and UCBMN-derived erythropoiesis were continuous and dynamic. We next used Monocle (Qiu et al., 2017) to construct pseudo-time trajectories of erythroid differentiation from both sources and clarify the key driving factors in UCBMN and PSC erythropoiesis. The branch point position of the pseudo-time trajectories indicates that cells from both origins differentiated along distinct early and late paths (Figures 2A, S2A-B). We then compared the differentially expressed genes (DEGs) in the early and late paths of both pseudo-time trajectories, analyzed the expression of the top 20 DEGs in each path, and observed the dynamic changes in their expression patterns. We observed conservatively higher expression of several ribosomal proteins during the early path compared to the late path, indicating ribosome biogenesis is an indispensable event for two origins during ex vivo erythropoiesis at early stage; however, other conserved genes such as solute carrier family 25 member 37 (*SLC25A37)* and interferon-stimulated gene 15 (*ISG15)* exhibited divergent expression pattern in both pseudo-time trajectories (Figures 2B-C). Of these, *SLC25A37* could play a role in heme biosynthesis, at late path for UCBMN but early path for PSC, during ex vivo erythropoiesis.

**Figure. 2.**
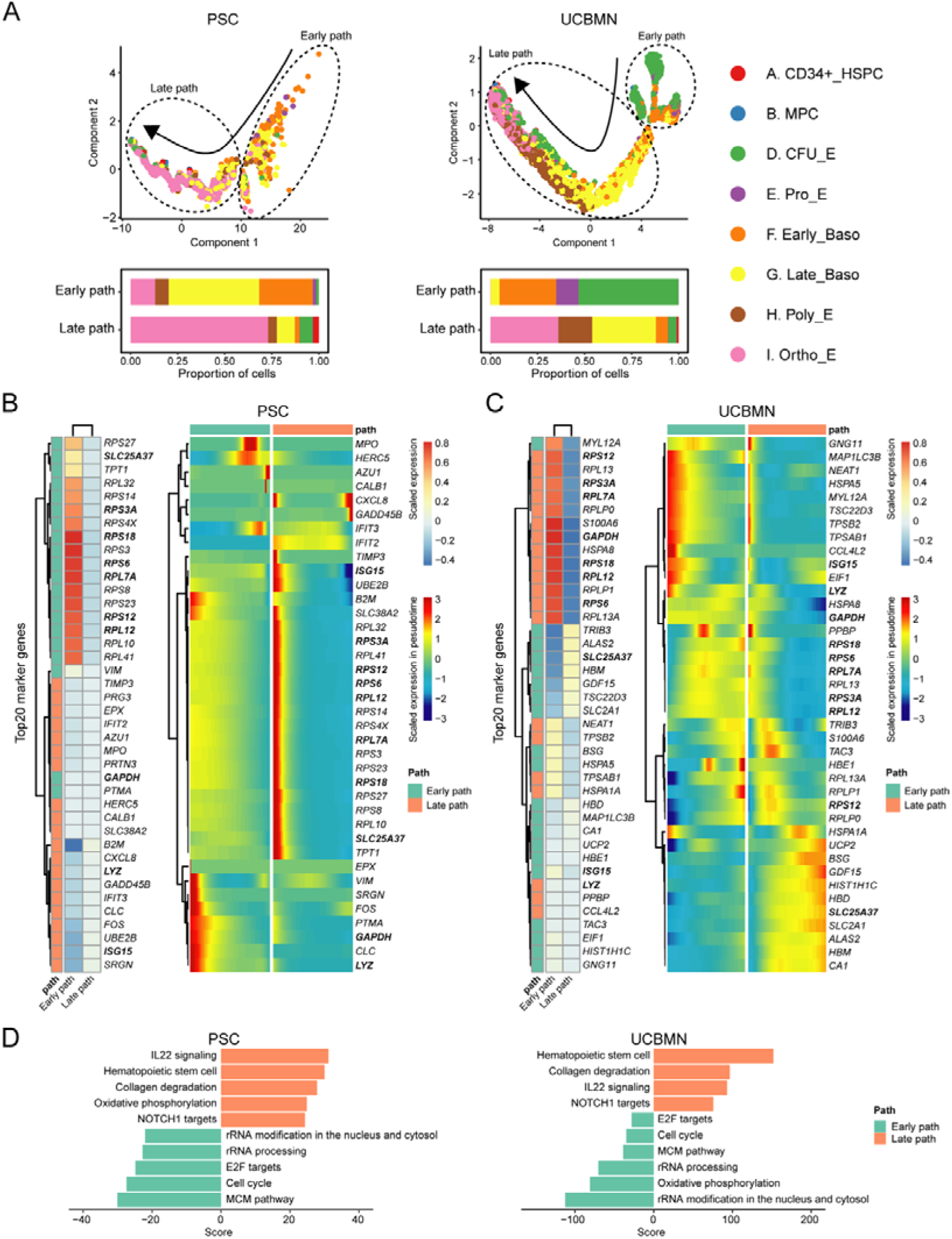
Regulatory dynamics of UCBMN- and PSC-derived erythropoiesis. (A) Developmental pseudo-time trajectories (top) of progenitor and erythroid cells and the proportions of cells (bottom) for PSC and UCBMN origin in early and late paths. Colors are indicating different cell types. BFU-E was not applied for this analysis because the subgroup with cell number less than 10 was removed in scRNA-Seq analysis. (B) Heatmap plots are representing the average scaled expression levels (left) and the pseudo-time expression levels (right) of top 20 marker genes (ordered by averaged log2 fold change, P value < 0.05) between the early and late paths in PSC derived progenitors and erythroid cells. (C) Heatmap plots are representing the average scaled expression levels (left) and the pseudo-time expression levels (right) of top 20 marker genes (ordered by averaged log2 fold change, P value < 0.05) between the early and late paths in UCBMN derived progenitors and erythroid cells. (D) Bar plot is showing GSVA enrichment scores of PSC (left) and UCBMN (right) derived cells in the early and late paths.

By trajectory analysis (Hao et al., 2021), we identified the expression levels of nuclear paraspeckle assembly transcript 1 (NEAT1) and eukaryotic translation initiation factor 1 (*EIF1*) in the early path, and the specific expression of stress erythropoiesis-related genes such as Tribbles pseudokinase 3 (*TRIB3)* and growth differentiation factor 15 (*GDF15)* (Dev et al., 2017; Hao et al., 2019) in the late path of UCBMN-derived erythropoiesis. The expression of globin gene hemoglobin subunit epsilon 1 (HBE1) occurred earlier in cell differentiation than that of hemoglobin subunit delta (HBD) and mu (HBM) (Fig. 2C). In the differentiation trajectory of PSC-derived erythroid cells, C-X-C motif chemokine ligand 8 (*CXCL8)* and growth arrest and DNA damage inducible beta (*GADD45B)* were specifically expressed in certain cells in the late path. We detected no globin-related genes among the top 20 DEGs of the PSC-derived cells (Figure 2C), indicating that UCBMN- and PSC-derived terminal erythroid differentiation are differentially regulated.

We performed gene set variation analysis (GSVA) (Hänzelmann et al., 2013), and identified several conserved functions, including rRNA processing, IL22 signaling, and the cell cycle, in erythroid differentiation from both sources (Figures 2D-E, S2C-E). However, the oxidative phosphorylation pathway was enriched in the early path of UCBMN-derived cells but not in the late path of the PSC-derived cells, suggesting a difference in energy demand during ex vivo erythropoiesis.

### Regulatory TF networks in Ortho-Es derived from *in vitro* erythropoiesis

Regulon activation was stage-specific during erythropoiesis. We next used SCENIC to illustrate regulons activation at different stages of erythropoiesis from both sources (Aibar et al., 2017). We observed dynamic and specific regulation at different developmental stages throughout erythropoiesis. The expression of certain regulons such as *RUNX1, KLF1, TAL1,* and *GATA1* was conserved during erythropoiesis, while other regulons such as *KLF3, NFE2,* and *MXI1* were specifically active in UCBMN-derived Ortho-Es (Figures 3A and S2F). This suggests that specific Ortho-Es regulons and the corresponding gene regulatory networks may be related to erythroid maturation.

**Figure. 3.**
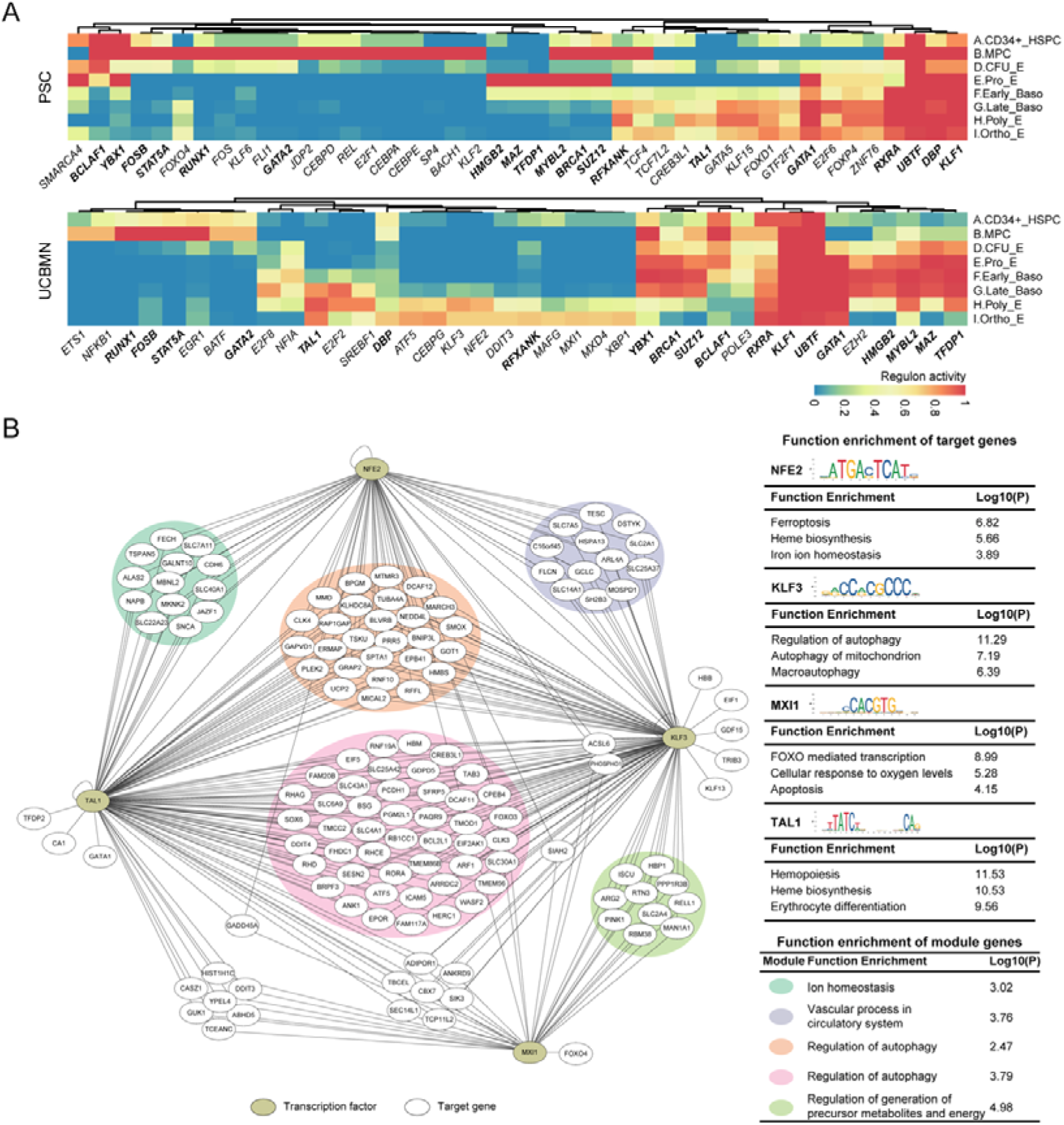
TFs dynamic networks involved in erythroid cell development and maturation in ortho-E. (A) Heatmap showing active regulon scores for each progenitor and erythroid cell type from PSC and UCBMN origin. (B) Networks of maturation-related TFs (olive color nodes) *NFE2*, *KLF3*, *MXI1*, and *TAL1* and their target genes (white nodes) in ortho-E. Function enrichment analysis of target genes and module genes are listed in tables. Network was constructed with Cytoscape v. 3.7.2 (https://cytoscape.org/download.html). Functional enrichment analysis was performed with Metascape (https://metascape.org/gp/index.html#/main/step1). TF sequences from JASPAR database (https://jaspar.genereg.net) shown here resemble those of the regulons analyzed.

To further investigate the functions of these active regulons in UCBMN-derived ortho-Es, we constructed interacting networks with the functional regulon target gene sets. In contrast to PSCs, we observed highly enriched autophagy, ferroptosis, ion homeostasis, and cellular response to oxygen levels in UCBMN-derived ortho-Es that is the stage closely related to the maturation of erythroid cells. It was previously reported that *KLF3* fine-tunes erythroid cell maturation by interacting with *KLF1* and its other family members(Ilsley et al., 2017). We found *KLF3* was enriched in autophagy that is implicated in erythroid maturation(Grosso et al., 2017), and regulated *HBB*, *EIF1*, *GDF15*, *TRIB3*, and *KLF13,* controlling hemoglobin expression and stress erythropoiesis (Figure 3B). Moreover, in ortho-E, we observed that NFE2 was related to ferroptosis and heme biosynthesis, *MXI1* was involved in FOXO-mediated transcription and cellular response to oxygen levels, and *TAL1* participated in hematopoiesis, heme biosynthesis, and erythrocyte differentiation (Fig. 3B). Overall, we identified an Ortho-E-specific regulon-associated network that may regulate the development and maturation of erythroid cells.

### Profiling comparison of hemoglobin expression between UCBMN and PSC origin

Globin expression is a critical issue in *in vitro* erythropoiesis. We extracted and compared globin gene expression in 16,702 progenitors and erythroid cells in nine different categories (CD34^+^ HSPC, MPC, BFU-E, CFU-E, Pro-E, early baso-E, late baso-E, poly-E, and ortho-Es) (Figure 4A). The expression of globin genes was gradually upregulated as erythropoiesis progressed (Figure 4B). UCBMN-derived erythroblasts mainly expressed adult- and fetal-type globin, whereas PSC-derived erythroblasts mainly expressed embryonic- and fetal-type globin (Figures 4A, S3A-C).

**Figure. 4.**
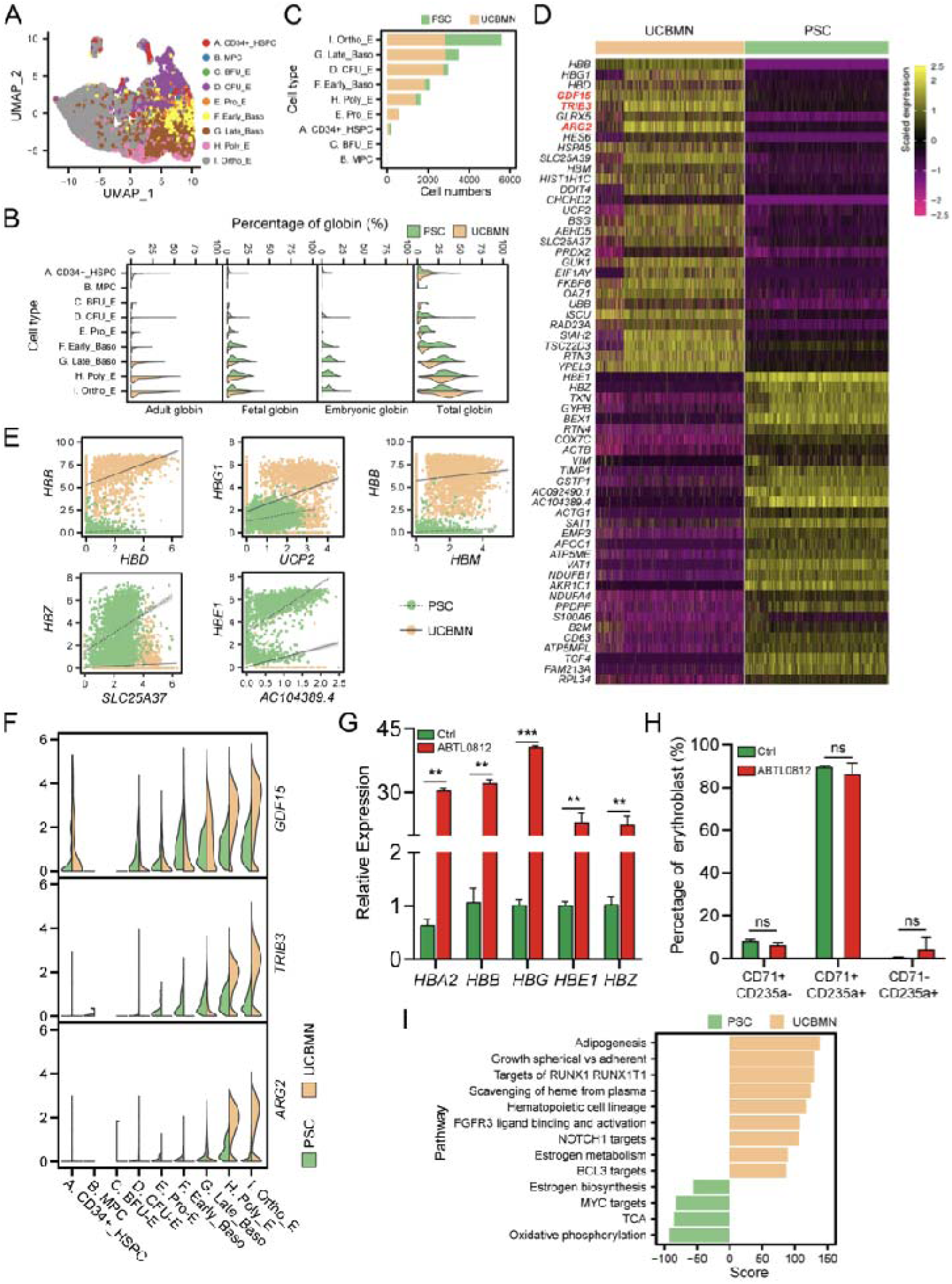
Profiling comparison of hemoglobin expression and the difference in stress erythropoiesis between UCBMN and PSC origin. (A) UMAP atlas of PSC- and UCBMN-derived progenitors and erythroid cells, respectively. Labels indicate cell type annotations performed by SingleR. (B) Violin plot showing % of adult, fetal, embryonic, and total globin for PSC-derived (green) and UCBMN-derived (yellow) progenitors and erythroid cells. (C) Bar plot comparing numbers of each cell type in proportions of cells differentiating towards erythroid cells derived from UCBMN and PSC origin (different colors). (D) Heatmap showing scaled average expression of top 30 most variable DEGs (in order of log2 FC) in PSC- and UCBMN-derived Ortho-E. Stress erythropoiesis-related genes were labelled in red. (E) Dot spot showing correlations between DEGs and globin genes in ortho-E derived from UCBMN and PSC origin. (F) Violin plot showing expression levels of *TRIB3*, *GDF15*, and *ARG2* in various cell populations derived from PSC (green) and UCBMN (yellow). (G) Bar plot showing significant (p<0.001) increase in hemoglobin expression relative to *18S* after treatment with specific *TRIB3* agonist ABTL-0812. **: P < 0.01, ***: P < 0.001. UCB-E were treated with 10μM ABTL-0812 from day 10, then perform rt-qPCR at day 22 (n=3). (H) Flow cytometry showing increase in CD71^-^CD235a^+^ after treatment with *TRIB3* agonist ABTL-0812 in (H). (I) Representative GSVA enrichment terms of DEGs for ortho-E derived from UCBMN and PSC origin.

To assess hemoglobin expression in PSC-derived and UCBMN-derived erythropoiesis, we compared the DEGs in ortho-Es between both origins. We found that ortho-E cell numbers were relatively equal, unlike the other erythroid cell developmental stages (Figures 4C-D, S3C). Spearman’s coefficients between the top 30 DEGs in the ortho-Es from both sources were used to identify genes with the highest correlations to the globin-encoding gene (Figures 4E, S3D-E), suggesting that these genes could participate in the expression of globin genes during ex vivo erythropoiesis from two origins, of which *UCP2* and *SLC25A37* are mitochondrial proteins, with *SLC25A37* playing an essential role in heme biosynthesis(Chen et al., 2009).

Among the top 30 DEGs, we observed that *GDF15*, *TRIB3*, and arginase 2 (*ARG2)* were related to stress erythropoiesis and were significantly upregulated in UCBMN-derived ortho-Es but not PSC (Figure 4F), indicating that the lack of β-globin in PSC-E may attribute to their deficiencies in stress erythropoiesis regulation. We evaluated the function of *TRIB3* in stress erythropoiesis and found that ABTL-0812, an TRIB3 agonist, leads to TRIB3 overexpression in UCB-E and significantly increased hemoglobin production, which is also revealed by the increase in CD71^-^CD235^+^ cells during terminal erythropoiesis (Figures 4H-I).

Moreover, we characterized ortho-E by performing gene set variation analysis (GSVA) and GO functional enrichment analysis of the top 30 DEGs between the two origin groups. We observed that ortho-Es from both sources had similar functions including structural ribosome constituent formation, rRNA binding, and peroxidase biosynthesis (Figures S3D). However, oxygen carrier activity was specifically enriched in UCBMN-derived ortho-Es, in addition to heme scavenging from plasma and estrogen metabolism (Figures 4G, S3D). By contrast, pathways related to the tricarboxylic acid cycle (TCA) and estrogen biosynthesis were significantly enriched in PSC-derived ortho-Es (Figures 4G).

### PSC-derived erythroid cells showed impaired expansion due to an abnormal cell cycle

We used the scran package in R to compare cell cycle ratios during erythroid differentiation between the two source groups (Lun et al., 2016) (Figure 5A). We observed that UCB-derived erythroid cells retained higher proliferation capacity than PSC as the pseudo-time trajectory progressed, because cell cycle analysis revealed that > 95% of all PSC-derived cells were in G1 and > 10% of all UCB-derived cells were in S (Figure 5B, C). The *RUNX1* regulon was upregulated in UCB progenitor cells. However, most of them lacked proliferative capacity (Figures 3A, 5C). We proposed that constitutive upregulation of *RUNX1* may explain why some progenitors cannot be induced into certain erythroid cells. Cell cycle regulation is a critical process in erythroid cell proliferation. An EdU FAC analysis between UCB-E D21 and PSC-E D18 indicated that the UCBMN-derived erythroid cells retained continuous proliferation abilities (Figure 5D).

**Figure. 5.**
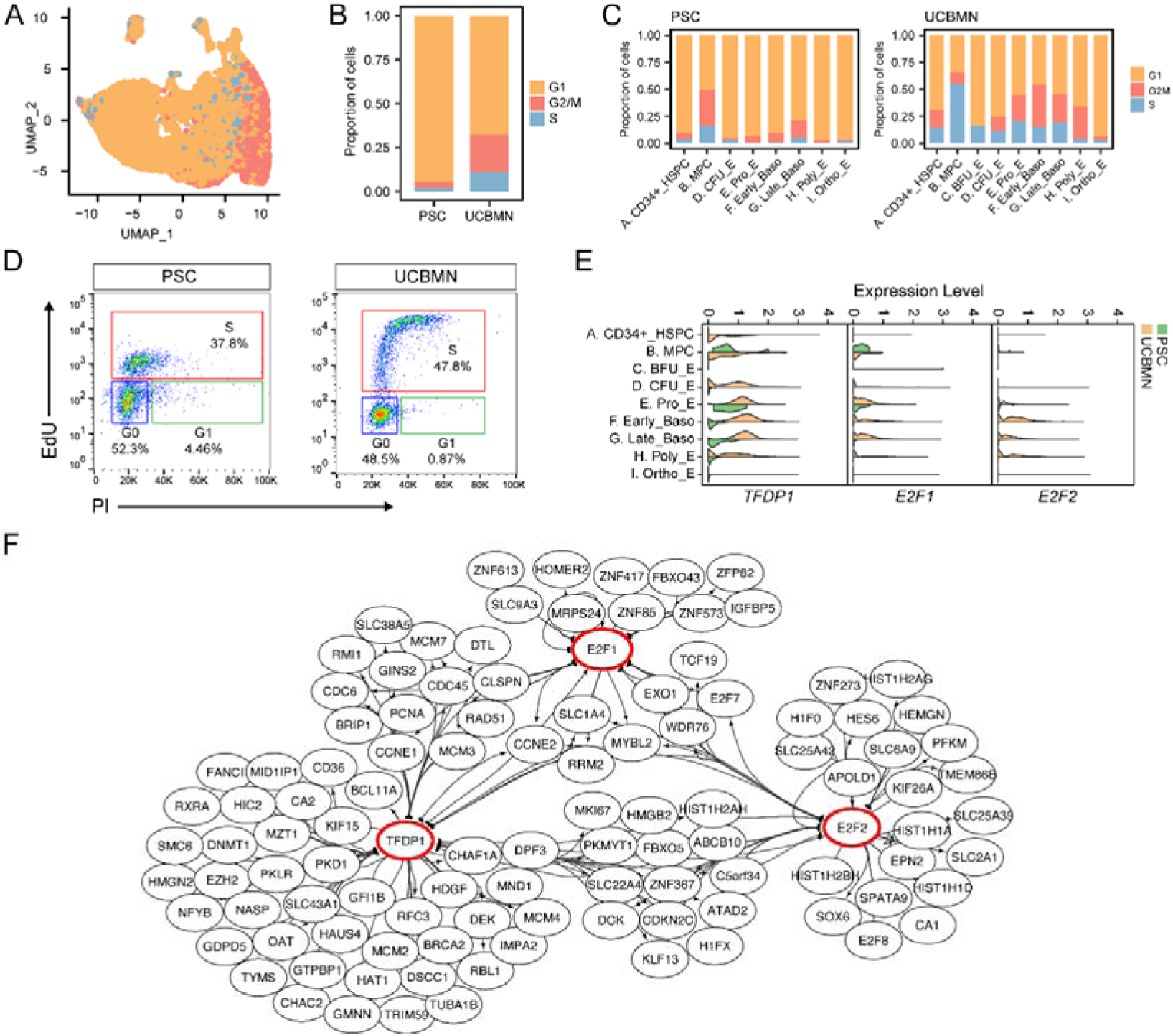
Cell-cycle states of PSC- and UCBMN-derived erythroid cells. (A) UMAP atlas of PSC- and UCBMN-derived progenitors and erythroid cells labeled with cell cycle annotation results from SingleR. (B) Bar plot showing proportions of cells derived from UCBMN and PSC origin in (A). (C) Bar plot showing proportions of each cell type in (A). (D) Flow cytometry analysis of cell cycle progression. DNA synthesis measured by EdU incorporation in D21 and D23 of UCB- and PSC-derived erythroid cells. (E) Violin plot showing *TFDP1*, *E2F1*, and *E2F2* TF expression levels in different cell populations derived from PSC (green) and UCBMN (yellow). (F) Network analysis of cell cycle-related *TFDP1*, *E2F1*, and *E2F2* TFs (red cycle nodes) and their target genes. Arrow direction indicate TFs targeted to genes. Network was plotted with Cytoscape v. 3.7.2.

*TFDP1* is a key cell cycle regulator during erythropoiesis(Li *et al*., 2014). In UCBMN-derived cells, *TFDP1* was active during various stages of development, from CFU-E to poly-E. However, in PSC-derived cells, it was expressed only in pro-E (Figures 5E). Cyclin-dependent kinases (*CDK*s) (Wang *et al*., 2022) and *TFDP1* are implicated in G2/S transformation. Relative differences in the activity levels of *TFDP1*, E2F transcription factor 1 (*E2F1)*, and E2F2 regulons may account for the reduced proliferation during erythropoiesis *in vitro* (Figures 5E). The target gene network comprising *TFDP1*, *E2F1*, and *E2F2* enriched the proliferation Ki-67 (*MKi67)* marker, CDK kinase gene (*CDKN2C)*, DNA replication initiator genes cell division cycle 6 (*CDC6)*, and *CDC45*. *TFDP1* targeted the *CD36* erythroid gene and BAF chromatin remodeling complex subunit (*BCL11A)* (Figure 5F) hemoglobin switch gene, indicating that *TFDP1* may regulate erythroid cell proliferation and promote erythroid differentiation.

### CD99^high^ progenitor cells represent a novel subpopulation of cells with higher proliferation ability

We are aware that relatively high percentages of CFU-E but not BFU-E are present in terminally differentiated cells, which limits the expansion capacity of in vitro erythropoiesis. FAC and CFU assay also detected a low percentage of CFU-E during terminal erythropoiesis derived from both origin (data not shown). We analyzed transcript expression in CFU-E and HSPC developmental trajectories using Monocle. This result revealed strong subclusters of new progenitor cells in CFU-E (Figure 6A). HSPC developed into early and late CFU-E along the erythropoiesis path (Figures 6A, S4A-S4I). A GO enrichment analysis showed that early CFU-E were enriched mainly in location maintenance and myeloid differentiation, indicating that early CFU-E corresponded to hematopoietic progenitor cells. However, late CFU-E were enriched in oxidative phosphorylation and oxygen transport, which correspond to definitive erythroid progenitor cells (Figures 6B-C, S4L, S4J). A cell cycle analysis showed that 25% and 6% of the cells in early and late CFU-E, respectively, were in phase S (Figure S4K).

**Figure. 6.**
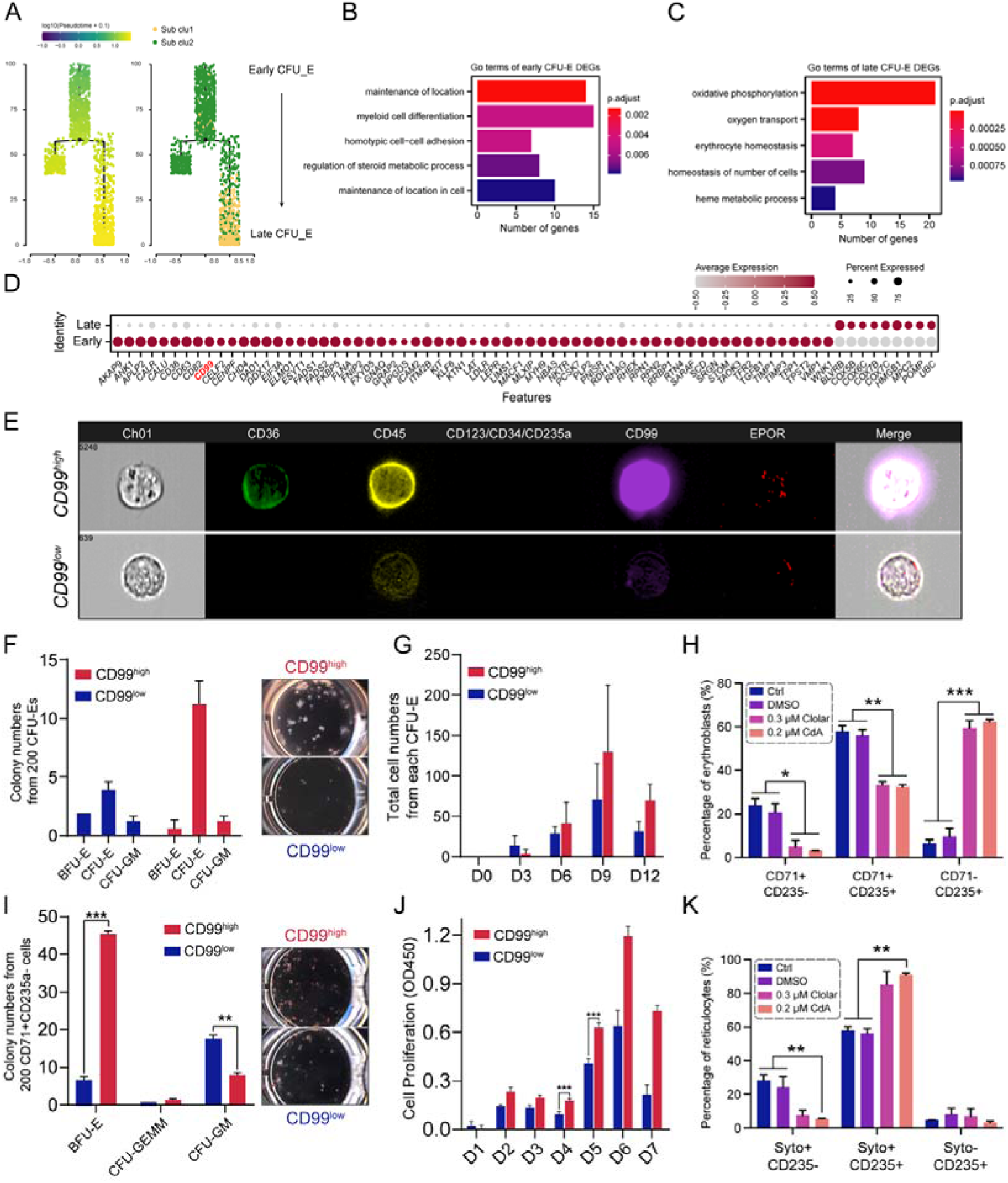
CD99^high^ population generates more erythroid cells than CD99^low^ population in erythroid progenitor cells. (A) Developmental pseudo-time trajectory of CFU-E cells labeled by pseudo-time scores (left) and CFU-E subsets (right). Arrow indicates the direction of differentiation. (B-C) Bar plot representing GO term enrichment results for DEGs between early (B) and late (C) CFU-E subsets. Bar colors indicate adjusted P-values of GO terms. (D) Dot plot showing membrane-encoding DEGs between early and late CFU-E cells. Colors indicate scaled average gene expression levels. Dot size indicates % of cells expressing genes in early and late CFU-E subsets. Membrane-endoding genes were obtained from UniProt under the key words membrane protein AND organism:“Homo sapiens (Human) [9606] ”, then overlapped the DEGs between early and late CFU-E cells to achieve the genes list as presented. (E) Image flow analysis of CD99 expression in UCB-generated CFU-E (CD235a^-^CD123^-^CD71^+^CD34^-^CD36^+^) cells. Image of cells was captured by Amnis ImageStreamX. (F) Bar plot of numbers and representative pictures of colonies formed by UCBMN derived CFU-E subpopulations in H4434 medium (n=3). (G) Bar plot of numbers of cells formed by single CFU-E subpopulation cells. CD99^high^ subpopulation generated two fold cells than CD99^low^ one, even no significant difference were detected between the two groups (n=3). (H) Bar plot comparing the percentage of erythroblasts treated with antagonist of CD99. UCBMN derived cells were treated with antagonist of CD99, 0.3µm Clolar and 0.2µm 2-CdA, from D10 (n=3). *: P < 0.5, **: P < 0.01, ***: P < 0.001. (I) Bar plot of numbers and representative pictures of colonies formed by UCB-CD34^+^ cells derived CD235a^-^CD71^+^ subpopulations in H4434 medium (n=4). **: P < 0.01, ***: P < 0.001. (J) The proliferation potent of UCBMN derived CFU-E subpopulation were tested by OD450 with CCK8 kit (n=3). ***: P < 0.001. (K) Bar plot of flow cytometry analyzed enucleated reticulocytes(Syto^-^CD235a^+^) percentage of (H). **: P < 0.01.

CD99 is a surface glycoprotein related to T cell migration and acute carcinoma progression (Angelini et al., 2015; Mannion et al., 2021) (Feng et al., 2021; Vaikari et al., 2018). In BM CD34^+^ cells, the CD99^high^ population has a higher migration potential than the CD99^low^ population (Imbert et al., 2006). Among the most highly variable genes between the two subclusters, we selected *CD99* to distinguish between early and late progenitor cells (Figure 6D). CFU-E subpopulations were isolated from UCBMN-derived cells as previously reported (Li *et al*., 2014) (Figure S4M). Giemsa staining revealed that CD99^high^ cells had larger nuclei than CD99^low^ cells. The latter occurred mainly among early progenitor cells (Figure S4N). Flow imaging and FACS analysis of the hematopoietic progenitor cells (HPCs) revealed that CD99 expression levels in the CFU-E were consistent with the results of the transcriptome analysis (Figures 6E, S6D).

By integrating the human bone marrow data (Jason D Buenrostro, 2018), we confirmed that early and late CFU-E cells in vivo were also marked by CD99^high^ and CD99^low^ (Figures S5A-D). The GO terms showing DEGs enriched in metabolic pathways indicate that the CD99^high^ subpopulation was enriched in HPC differentiation-related hemopoiesis regulation, whereas the CD99^low^ subpopulation was enriched in erythrocyte maturation-associated chromosome condensation. Thus, the CD99^high^ subcluster was predicted to have high *in vivo* proliferation capacity (Figure S5E).

We analyzed the colony-forming units and proliferation potential of both populations to evaluate the functional differences between the CD99^high^ and CD99^low^ progenitor cells. CD99^high^ progenitor cells generated larger and more numerous colonies than CD99^low^ progenitor cells (Figure 6F). CD99^high^ cells generated twice as many erythroblasts as CD99^low^ cells (Figure 6G). We verified these trends in CD235a^-^CD71^+^ cells using CFU and CCK8 assays (Figures 6H-I). More importantly, we observed increased production of mature erythroblasts (CD235a+/CD71-) and reticulocytes when CD99 agonists clofarabine (Clolar) and 2-chlorodeoxyadenosine (2-CdA) (Figures 6J-K) were used. CD99 agonists can be used to increase erythrocyte production in vitro.

### Macrophages were involved in *in vitro* erythropoiesis by cell-cell contact

As illustrated in Figure 1A, appropriately 10% of cells were double-negative for CD71/CD235a after terminal erythroid differentiation. At that point, the most significant non-erythroid cells that were in contact with erythroid cells were macrophages (Figure S6A). Macrophage-CFU-E communication decreased as the latter developed (Figure S7A). We used cell-cell contact and cell signaling databases in CellChat (version 1.1.2) of R to predict the ligand-receptor interactions between macrophages and erythroid cells. We identified a total of 30 and 36 ligand-receptor interaction pairs from macrophages to erythroid cells and erythroid cells to macrophages, respectively. Communication from macrophages to erythroid cells is more significant than the opposite (Figure S6B). CFU-E in our data were divided into early and late CFU-E cells with different *CD99* expression. Analysis of the communication between CFU-E subpopulations and macrophages showed that signaling and contact decreased from early to late stage erythropoiesis (Figure S7A). Furthermore, CD99-CD99 was the most significant communication ligand-receptor pair (Figures S7B-C) and the expression of *CD99* in CFU-E population also decreased as development progressed (Figures S7D-E). Our results demonstrated that macrophages are involved during in vitro erythropoiesis based on *CD99* expression. Specialized CFU-E cells interact with macrophages by establishing a CD99-CD99 contact.

## DISCUSSION

To clarify the mechanisms underlying the known limitations of RBC maturation and production ex vivo, we decoded the single-cell transcriptomics of UCBMN- and PSC-derived terminal erythroid cells. At the sequencing timepoint, morphology analysis of cells generated from two resources showed that the cells reached terminal stage of erythropoiesis (Figure 1A) ; Appropriately 90% of cells collected from both sources entered erythropoiesis, while 10% of cells remained both CD71 and CD235a negative, including non-erythroid cells and quiescent progenitors, which have been not previously described. Some hematopoietic and erythroid progenitor cells, such as CD34^+^ cells, BFU-E, and CFU-E, remained in an undifferentiated state, which might result from the heterogeneity, epigenetic modifications, or metabolic programming of the origin cells. We noted the diversity in the terminally differentiated cells, and the cell populations originating from the two sources were distributed in various development stages, which was consistent with the continuous differentiation model. Based on the cell-cell interaction analysis, in addition to macrophages, non-erythroid cells, including myelocytes, monocytes, neutrophils, and fibroblasts could also be involved in and facilitate in vitro erythropoiesis.

Macrophages participate in erythroid differentiation as a microenvironment component and contact erythroid cells via specific protein binding (Jacobsen et al., 2015; Javan et al., 2018). We observed marked differences in macrophage proportions in terminally differentiated cells from both sources, suggesting that macrophages were distinct, probably differentiated unequally, and performed different roles for each source. Besides erythroid cells, macrophages also facilitate the expansion of hematopoietic cells. Gao et al. reported that macrophages facilitate the formation of functional HSC/MPP units in the fetal liver to promote HSC expansion through growth factor secretion (Gao et al., 2022), and CD34 cell accumulation is increased after co-culture with M2-MΦs, but not M1-MΦs (Luo et al., 2018). Application of IL-6 in the absence of IL-1β is a vital driver for the differentiation of myeloid precursors and monocytes(Mantovani et al., 2019; Mitroulis et al., 2018). We hypothesized that by increasing the proportion of alternatively-polarized macrophages and moderately supplementing the medium with IL-1β and IL-6 during the proliferation stage, we could improve *in vitro* erythropoiesis. In the current study, we observed the interactions between macrophages and CFU-E during ex vivo erythropoiesis through the CD99-CD99 pathway, which may be a novel mechanism by which macrophages participate in erythropoiesis.

We observed that stress erythropoiesis specifically occurred in UCBMN-derived ortho-Es, and related genes including *TRIB3*, *ARG2*, and *GDF15* were upregulated. *TRIB3* strongly responded to insufficient erythropoietin (EPO) in adult BM EPCs (Sathyanarayana et al., 2008). *TRIB3^-/-^* mice had elevated RBC counts and hemoglobin content. These studies demonstrate that *TRIB3* regulates stress erythropoiesis (Dev *et al*., 2017). The current study firstly identify the small molecule ABTL-0812, α-hydroxylinoleic acid, as an agent that increases hemoglobin expression and promotes erythroid maturation. *GDF15* regulates progenitor metabolism and promotes stress erythropoiesis in mouse models (Hao *et al*., 2019). *ARG2* confers immunosuppressive properties and is associated with the development of the neonatal immune system (Elahi and Mashhouri, 2020), and remains to be evaluated in ex vivo erythropoiesis. In humans, the levels of CD71^+^ erythroid progenitor cells were increased in spleens when stress erythropoiesis occurred (Elahi, 2019). Culturing HSPCs in medium containing stem cell growth factor (SCF), EPO, and dexamethasone mimics stress erythropoiesis (Heshusius et al., 2019; Hsieh et al., 2022). We hypothesized that UCB-derived progenitors could resemble stress erythroid progenitors, and UCB-E-enriched transcripts *TRIB3*, *ARG2*, and *GDF15* were potential regulators of hemoglobin expression. The present work showed less immune cells in UCB-E, such as eosinophils and neutrophils, than PSC-E, which could be related to stress erythropoiesis. Further understanding of the mechanisms behind the *in vitro* stress erythropoiesis will aid erythrocyte regeneration efforts.

This is the first study, to the best of our knowledge, to identify CD99 as a marker of early and late CFU-E in UCB-E and PSC-E, as well as in BM. Furthermore, the CD99^high^ subpopulation in CFU-E was a subcluster with high proliferative capacity both *in vitro* and *in vivo*. Identification of this subpopulation in the exponential growth period of erythropoiesis, from BFU-E to CFU-E, provided new insights on the mechanism of erythroid progenitor proliferation. It has been shown that CD99 is involved in cell expansion through the HIF1a-CD99-ERK1/2 pathway (Feng *et al*., 2021). In this study, we evaluated CD99 expression and observed that it was expressed in the early stages of erythropoiesis, and decreased gradually during erythropoiesis. Reducing CD99 expression using antagonists facilitated erythroid maturation, shrinking the population of undifferentiated cells in the culture system. This finding provides new knowledge on the role of CD99 in erythroid progenitor proliferation and maturation during ex vivo erythropoiesis.

In this study, we comprehensively compared UCB- and PSC-derived terminally differentiated erythroid cells at the single cell level and elucidated the mechanisms underlying current limitations of *in vitro* erythropoiesis. For the first time, we deciphered the cell composition and differentiating path, and determined regulons potentially contributing to maturation, PSC-E proliferation, and globin expression by comparing distinct cells at the single cell level. We observed the heterogeneity of CFU-E that was divided by CD99 expression. We also identified a novel subpopulation with high proliferation capacity in erythroid progenitors, and its putative role in ex vivo RBC generation, as well as small molecules which increase hemoglobin expression that can be applied to future RBC regeneration efforts. Future research should endeavor to understand the specific effects of these regulators on erythropoiesis and determine how they can be applied to improve RBC production. These investigations will serve as a reference for potential large-scale, in-depth research on RBC generation.

## ACKNOWLEDGMENTS

This research was supported by the Strategic Priority Research Program of the Chinese Academy of Sciences (No. XDA16010602 to X.F.), National Natural Science Foundation of China (No. 81870097 to Z.Z, No. 82070114 and 82270126 to X.F.), the National Key Research and Development Program of China (No. 2017YFA0103100, No. 2017YFA0103103 and No. 2017YFA0103104 to X.P.), and Science and Technology Program of Guangzhou, China (No. 202002030025).

## AUTHOR CONTRIBUTIONS

X.W. and W.Z. performed research and analyzed the data; X.W. collected, and interpreted data with help of H.Y., T.C., R.Z., L.Z., and X.L.; W.Z. performed bioinformation analysis with help of S.Z. and Z.X.; X.P., X.F., Z. Z, W.Y., J.X., X.W., W.Z., Z.Z designed research, and wrote the manuscript; X.P., X.F., W.Y., and Z. Z supervised and funded the project; and all authors have read and approved the final manuscript.

## DATA AVAILABILITY

The scRNA-seq datasets were available in Genome Sequence Archive for Human (https://ngdc.cncb.ac.cn/gsa-human/s/AdGSYf73). This paper does not report original code. Any additional information required to reanalyze the data reported in this paper is available from the lead contact upon request.

## STAR METHODS

### Key resources table

| REAGENT or RESOURCE | SOURCE | IDENTIFIER |
| --- | --- | --- |
| **Antibodies** |  |  |
| CD235a-PEcy7 | Biolegend | 349112 |
| CD123-PEcy7 | BD Biosciences | 560826 |
| CD71-BV605 | BD Biosciences | 743306 |
| CD34-PE | Biolegend | 343506 |
| CD36-APC | BD Biosciences | 550956 |
| CD99-BV421 | BD Biosciences | 743040 |
| CD45-BV786 | BD Biosciences | 563716 |
| CD34-PEcy7 | BD Biosciences | 560070 |
| CD36-PE | Biolegend | 336026 |
| EPOR-APC | R&D Systems | FAB3072A |
| CD71-APC | BD Biosciences | 551374 |
| CD71-PEcy7 | Biolegend | 334112 |
| CD235a-BV421 | BD Biosciences | 562938 |
| Brilliant Stain Buffer | BD Biosciences | 563794 |
| Syto16 | Invitrogen | S7578 |
| **Biological samples** |  |  |
| Human umbilical cord blood | Umbilical Cord Blood Bank, Beijing, China | AF/SC-08/02.177 |
| **Chemicals, peptides, and recombinant proteins** |  |  |
| EdU (5-Ethynyl-2-deoxyuridine) | Beyotime | C0078S |
| TRIzol Reagent | Invitrogen | 15596026 |
| Clofarabin (Clolar) | Selleck | S1218 |
| Cladribine (RWJ26251, CdA) | Selleck | S1199 |
| ABTL-0812 (α-Hydroxylinoleic acid) | Selleck | S9611 |
| StemRegenin 1 (SR1, aryl hydrocarbon receptor antagonist) | STEMCELL | 72342 |
| Benzidine | Chemical Book | 92-87-5 |
| Recombinant human FGF basic | PeproTech | 100-18C |
| Recombinant human SCF | PeproTech | 300-07 |
| Recombinant human BPM4 | R&D Systems | 314-BP |
| Recombinant human VEGF 165 | R&D Systems | 293-VE |
| Recombinant human TPO | PeproTech | 300-18 |
| Recombinant human IL3 | PeproTech | 300-03 |
| Recombinant human IL6 | PeproTech | 300-06 |
| Recombinant human EPO | PeproTech | 100-64 |
| Recombinant human Flt3-Ligand | PeproTech | 300-19 |
| Recombinant human IGF1 | PeproTech | 100-11 |
| Critical commercial assays |  |  |
| Giemsa staining kit | BASO | BA4017 |
| Deposited data |  |  |
| scRNA-Seq | This paper | GEO: GSE166426 |
| Experimental models: Cell lines |  |  |
| Runx1C-H9 | Dan Kaufman | University of California (San Diego) |
| Oligonucleotides |  |  |
| Primers for rt-qPCR, see Table S1 | This paper | N/A |
| Software and algorithms |  |  |
| R version 4.0.3 | http://www.r-project.org/ | N/A |
| Prism version 9.0.0 | GraphPad | N/A |
| Flowjo version 10 | https://www.flowjo.com | N/A |
| Cytoscape version 3.7.2 | https://cytoscape.org/download.html | N/A |

## EXPERIMENTAL MODEL AND SUBJECT DETAILS

### Samples

Human umbilical cord blood was acquired from the Umbilical Cord Blood Bank, Beijing, China (AF/SC-08/02.177). Human Runx1C-H9 cells were provided courtesy of Prof. Dan Kaufman of the University of California (San Diego), San Diego, CA, USA. We applied single cell RNA sequencing on days 21 and 23 of UCB-E and PSC-E.

### Generation of erythrocyte from UCBMN

A four stages protocol of production erythrocytes from UCBMNs were modified from Qin et al. described before[9]. Stage 1 consisted of enrichment from days 0-7. The cells were cultured with 1 × 10^6^/mL SFEM II (STEMCELL Technologies, Vancouver, BC, Canada), 100 ng/mL rhSCF (PeproTech, Rocky Hill, NJ, USA), 50 ng/mL rhIL3 (PeproTech), 30 ng/mL rhIL6 (PeproTech), and 10 ng/mL rhTPO (PeproTech). Stage 2 consisted of proliferation from days 8-14. The cells were cultured with 5 × 10^5^/mL SFEM II (STEMCELL Technologies), 100 ng/mL rhSCF (PeproTech), 50 ng/mL rhIL3 (PeproTech), and 3 IU/mL rhEPO (PeproTech). Stage 3 consisted of differentiation from days 15-21. The cells were cultured in 5 × 10^5^/mL SCGM (CellGenix, Freiburg im Breisgau, Germany), 50 ng/mL rhSCF (PeproTech), 40 ng/mL rhIGF1 (PeproTech), 3 IU/mL rhEPO (PeproTech), 1×lipid (Gibco; Thermo Fisher Scientific, Waltham, MA, USA), and 200 ng/mL transferrin (Sigma-Aldrich). Stage 4 consisted of maturation from days 21-28. The cells were cultured in 5 × 10^5^/mL SCGM (CellGenix), 3 IU/mL rhEPO (PeproTech), 200× P188 (Gibco), and 200 ng/mL transferrin (Sigma-Aldrich, USA).

### Erythrocyte generation from PSCs

PSCs were co-cultured with MEF layer cells in DEME/F12 (Gibco), 20% KSR (Gibco) supplemented with GlutaMax (Gibco), NEAA (Gibco), and 10 ng/mL bFGF (PeproTech). The PSCs were dissolved in TrypleSelect (Gibco) after reaching 80% confluence. From days 0-14, Spin EB was performed to induce the PSCs to the mesoblastema stage. The hematopoietic stem cells were collected and the progenitor cells emerged from EB [10]. From day 15 onwards, the PSC-derived HSPCs were cultured in the same manner as the Stages 3-4 UCBMNs.

### scRNA-Seq

The unsorted cells were processed for library preparation with a Chromium Single Cell 3’ Reagent Kit v. 3 (10x Genomics, USA). Indexed libraries were pooled and sequenced on a NovaSeq 6000 platform (Illumina, USA) using 150-bp paired-end reads.

### SCENIC analysis

UMI count matrix data obtained by Seurat were used as the input for the SCENIC v. 1.2.4 package in R (R Core Team, Vienna, Austria) to predict changes in the active TFs during erythropoiesis[11]. The cisTarget Human motif database v. 9 (https://resources.aertslab.org/cistarget/motif2tf/motifs-v9-nr.hgnc-m0.001-o0.0.tbl) comprising 24,453 motifs was used with its default settings to enrich the gene signatures and isolate targets from them based on *cis*-regulatory cues. The “aucell” positional argument was used to detect regulon enrichment across single cells. The SCENIC v. 1.2.4 package in R loaded the “binary matrix” result and showed activated regulons. A heatmap was plotted with the pheatmap v. 1.0.12 module in R. The “regulon targets info” was loaded into Cytoscape v. 3.7.2 (https://cytoscape.org/download.html) to construct a network of TFs and their target genes.

### Quantification and statistical analysis

All data analysis was performed in R (version 4.0.3).

## REFERENCE

Aibar, S., Gonzalez-Blas, C.B., Moerman, T., Huynh-Thu, V.A., Imrichova, H., Hulselmans, G., Rambow, F., Marine, J.C., Geurts, P., Aerts, J., et al. (2017). SCENIC: single-cell regulatory network inference and clustering. Nat Methods 14, 1083–1086. 10.1038/nmeth.4463.

Angelini, D.F., Ottone, T., Guerrera, G., Lavorgna, S., Cittadini, M., Buccisano, F., De Bardi, M., Gargano, F., Maurillo, L., Divona, M., et al. (2015). A Leukemia-Associated CD34/CD123/CD25/CD99+ Immunophenotype Identifies FLT3-Mutated Clones in Acute Myeloid Leukemia. Clin Cancer Res 21, 3977–3985. 10.1158/1078-0432.Ccr-14-3186.

Aran, D., Looney, A.P., Liu, L., Wu, E., Fong, V., Hsu, A., Chak, S., Naikawadi, R.P., Wolters, P.J., Abate, A.R., et al. (2019). Reference-based analysis of lung single-cell sequencing reveals a transitional profibrotic macrophage. Nat Immunol 20, 163–172. 10.1038/s41590-018-0276-y.

Buenrostro, J.D., Corces, M.R., Lareau, C.A., Wu, B., Schep, A.N., Aryee, M.J., Majeti, R., Chang, H.Y., and Greenleaf, W.J. (2018). Integrated Single-Cell Analysis Maps the Continuous Regulatory Landscape of Human Hematopoietic Differentiation. Cell 173, 1535–1548. 10.1016/j.cell.2018.03.074.

Chen, W., Paradkar, P.N., Li, L., Pierce, E.L., Langer, N.B., Takahashi-Makise, N., Hyde, B.B., Shirihai, O.S., Ward, D.M., Kaplan, J., and Paw, B.H. (2009). Abcb10 physically interacts with mitoferrin-1 (Slc25a37) to enhance its stability and function in the erythroid mitochondria. Proc. Natl. Acad. Sci. U. S. A. 106, 16263–16268. 10.1073/pnas.0904519106.

Dev, A., Asch, R., Jachimowicz, E., Rainville, N., Johnson, A., Greenfest-Allen, E., and Wojchowski, D.M. (2017). Governing roles for Trib3 pseudokinase during stress erythropoiesis. Exp Hematol 49, 48–55. 10.1016/j.exphem.2016.12.010.

Elahi, S. (2019). Neglected Cells: Immunomodulatory Roles of CD71(+) Erythroid Cells. Trends Immunol 40, 181–185. 10.1016/j.it.2019.01.003.

Elahi, S., and Mashhouri, S. (2020). Immunological consequences of extramedullary erythropoiesis: immunoregulatory functions of CD71(+) erythroid cells. Haematologica 105, 1478–1483. 10.3324/haematol.2019.243063.

Feng, X.D., Zhu, J.Q., Zhou, J.H., Lin, F.Y., Feng, B., Shi, X.W., Pan, Q.L., Yu, J., Li, L.J., and Cao, H.C. (2021). Hypoxia-inducible factor-1α-mediated upregulation of CD99 promotes the proliferation of placental mesenchymal stem cells by regulating ERK1/2. World J Stem Cells 13, 317–330. 10.4252/wjsc.v13.i4.317.

Galat, Y., Elcheva, I., Dambaeva, S., Katukurundage, D., Beaman, K., Iannaccone, P.M., and Galat, V. (2018). Application of small molecule CHIR99021 leads to the loss of hemangioblast progenitor and increased hematopoiesis of human pluripotent stem cells. Experimental Hematology 65. 10.1016/j.exphem.2018.05.007.

Gao, S., Shi, Q., Zhang, Y., Liang, G., Kang, Z., Huang, B., Ma, D., Wang, L., Jiao, J., Fang, X., et al. (2022). Identification of HSC/MPP expansion units in fetal liver by single-cell spatiotemporal transcriptomics. Cell Res 32, 38–53. 10.1038/s41422-021-00540-7.

Grosso, R., Fader, C.M., and Colombo, M.I. (2017). Autophagy: A necessary event during erythropoiesis. Blood Reviews 31, 300–305. 10.1016/j.blre.2017.04.001.

Hänzelmann, S., Castelo, R., and Guinney, J. (2013). GSVA: gene set variation analysis for microarray and RNA-seq data. BMC Bioinf. 14, 1–15. 10.1186/1471-2105-14-7.

Hao, S., Xiang, J., Wu, D.C., Fraser, J.W., Ruan, B., Cai, J., Patterson, A.D., Lai, Z.C., and Paulson, R.F. (2019). Gdf15 regulates murine stress erythroid progenitor proliferation and the development of the stress erythropoiesis niche. Blood Adv 3, 2205–2217. 10.1182/bloodadvances.2019000375.

Hao, Y., Hao, S., Andersen-Nissen, E., Mauck, W.M., 3rd, Zheng, S., Butler, A., Lee, M.J., Wilk, A.J., Darby, C., Zager, M., et al. (2021). Integrated analysis of multimodal single-cell data. Cell 184, 3573–3587. 10.1016/j.cell.2021.04.048.

Heshusius, S., Heideveld, E., Burger, P., Thiel-Valkhof, M., Sellink, E., Varga, E., Ovchynnikova, E., Visser, A., Martens, J.H.A., von Lindern, M., and van den Akker, E. (2019). Large-scale in vitro production of red blood cells from human peripheral blood mononuclear cells. Blood Adv 3, 3337–3350. 10.1182/bloodadvances.2019000689.

Hsieh, H.-H., Yao, H., Ma, Y., Zhang, Y., Xiao, X., Stephens, H., Wajahat, N., Chung, S.S., Xu, L., Xu, J., et al. (2022). Epo-IGF1R crosstalk expands stress-specific progenitors in regenerative erythropoiesis and myeloproliferative neoplasm. Blood. 10.1182/blood.2022016741.

Hu, J., Liu, J., Xue, F., Halverson, G., Reid, M., Guo, A., Chen, L., Raza, A., Galili, N., Jaffray, J., et al. (2013). Isolation and functional characterization of human erythroblasts at distinct stages: implications for understanding of normal and disordered erythropoiesis in vivo. Blood 121, 3246–3253. 10.1182/blood-2013-01-476390.

Huang, P., Zhao, Y., Zhong, J., Zhang, X., Liu, Q., Qiu, X., Chen, S., Yan, H., Hillyer, C., Mohandas, N., et al. (2020). Putative regulators for the continuum of erythroid differentiation revealed by single-cell transcriptome of human BM and UCB cells. Proc. Natl. Acad. Sci. U. S. A. 117, 12868–12876. 10.1073/pnas.1915085117.

Ilsley, M.D., Gillinder, K.R., Magor, G.W., Huang, S., Bailey, T.L., Crossley, M., and Perkins, A.C. (2017). Krüppel-like factors compete for promoters and enhancers to fine-tune transcription. Nucleic Acids Res 45, 6572–6588. 10.1093/nar/gkx441.

Imbert, A.M., Belaaloui, G., Bardin, F., Tonnelle, C., Lopez, M., and Chabannon, C. (2006). CD99 expressed on human mobilized peripheral blood CD34+ cells is involved in transendothelial migration. Blood 108, 2578–2586. 10.1182/blood-2005-12-010827.

Jacobsen, R.N., Perkins, A.C., and Levesque, J.P. (2015). Macrophages and regulation of erythropoiesis. Curr Opin Hematol 22, 212–219. 10.1097/moh.0000000000000131.

Jacobsen, S.E.W., and Nerlov, C. (2019). Haematopoiesis in the era of advanced single-cell technologies. Nat Cell Biol 21, 2–8. 10.1038/s41556-018-0227-8.

Jason D Buenrostro, M.R.C., Caleb A Lareau, Beijing Wu, Alicia N Schep, Martin Aryee, Ravindra Majeti, Howard Y. Chang, and William J. Greenleaf (2018). Integrated single-cell analysis maps the continuous regulatory landscape of human hematopoietic differentiation. Cell 173, 1535–1548.

Javan, G.T., Salhotra, A., Finley, S.J., and Soni, S. (2018). Erythroblast macrophage protein (Emp): Past, present, and future. Eur J Haematol 100, 3–9. 10.1111/ejh.12983.

Li, J., Hale, J., Bhagia, P., Xue, F., Chen, L., Jaffray, J., Yan, H., Lane, J., Gallagher, P.G., Mohandas, N., et al. (2014). Isolation and transcriptome analyses of human erythroid progenitors: BFU-E and CFU-E. Blood 124, 3636–3645. 10.1182/blood-2014-07-588806.

Lun, A.T., McCarthy, D.J., and Marioni, J.C. (2016). A step-by-step workflow for low-level analysis of single-cell RNA-seq data with Bioconductor. F1000Res 5, 2122. 10.12688/f1000research.9501.2.

Luo, Y., Shao, L., Chang, J., Feng, W., Liu, Y., Cottler-Fox, M., Emanuel, P., Hauer-Jensen, M., Bernstein, I., Liu, L., et al. (2018). M1 and M2 macrophages differentially regulate hematopoietic stem cell self-renewal and ex vivo expansion. Blood Adv. 2, 859–870. 10.1182/bloodadvances.2018015685.

Mannion, A.J., Odell, A.F., Taylor, A., Jones, P.F., and Cook, G.P. (2021). Tumour cell CD99 regulates transendothelial migration via CDC42 and actin remodelling. J Cell Sci 134. 10.1242/jcs.240135.

Mantovani, A., Dinarello, C.A., Molgora, M., and Garlanda, C. (2019). Interleukin-1 and Related Cytokines in the Regulation of Inflammation and Immunity. Immunity 50, 778–795. 10.1016/j.immuni.2019.03.012.

Mitroulis, I., Ruppova, K., Wang, B., Chen, L.S., Grzybek, M., Grinenko, T., Eugster, A., Troullinaki, M., Palladini, A., Kourtzelis, I., et al. (2018). Modulation of Myelopoiesis Progenitors Is an Integral Component of Trained Immunity. Cell 172, 147–161. 10.1016/j.cell.2017.11.034.

Olivier, E.N., Marenah, L., McCahill, A., Condie, A., Cowan, S., and Mountford, J.C. (2016). High-Efficiency Serum-Free Feeder-Free Erythroid Differentiation of Human Pluripotent Stem Cells Using Small Molecules. Stem Cells Transl. Med. 5, 1394–1405. 10.5966/sctm.2015-0371.

Qiu, X., Mao, Q., Tang, Y., Wang, L., Chawla, R., Pliner, H.A., and Trapnell, C. (2017). Reversed graph embedding resolves complex single-cell trajectories. Nat Methods 14, 979–982. 10.1038/nmeth.4402.

Rallapalli, S., Guhathakurta, S., Narayan, S., Bishi, D.K., Balasubramanian, V., and Korrapati, P.S. (2019). Generation of clinical-grade red blood cells from human umbilical cord blood mononuclear cells. Cell Tissue Res 375, 437–449. 10.1007/s00441-018-2919-6.

Sathyanarayana, P., Dev, A., Fang, J., Houde, E., Bogacheva, O., Bogachev, O., Menon, M., Browne, S., Pradeep, A., Emerson, C., and Wojchowski, D.M. (2008). EPO receptor circuits for primary erythroblast survival. Blood 111, 5390–5399. 10.1182/blood-2007-10-119743.

Tao, L., Togarrati, P.P., Choi, K.-D., and Suknuntha, K. (2017). StemRegenin 1 selectively promotes expansion of Multipotent Hematopoietic Progenitors derived from Human Embryonic Stem Cells. J Stem Cells Regen Med 13, 75–79. 10.46582/jsrm.1302011.

Vaikari, V.P., Yang, J., Akhtari, M., and Alachkar, H. (2018). Functional Analysis of CD99 Upregulation in Acute Myeloid Leukemia. Blood 132, 5129–5129. 10.1182/blood-2018-99-120348.

Wang, L., Guan, X., Wang, H., Shen, B., Zhang, Y., Ren, Z., Ma, Y., Ding, X., and Jiang, Y. (2017). A small-molecule/cytokine combination enhances hematopoietic stem cell proliferation via inhibition of cell differentiation. Stem Cell Res. Ther. 8, 169. 10.1186/s13287-017-0625-z.

Wang, S., Zhao, H., Zhang, H., Gao, C., Guo, X., Chen, L., Lobo, C., Yazdanbakhsh, K., Zhang, S., and An, X. (2022). Analyses of erythropoiesis from embryonic stem cell-CD34(+) and cord blood-CD34(+) cells reveal mechanisms for defective expansion and enucleation of embryomic stem cell-erythroid cells. J Cell Mol Med 26, 2404–2416. 10.1111/jcmm.17263.

Xin, Z., Zhang, W., Gong, S., Zhu, J., Li, Y., Zhang, Z., and Fang, X. (2021). Mapping Human Pluripotent Stem Cell-derived Erythroid Differentiation by Single-cell Transcriptome Analysis. GPB 19, 358–376. 10.1016/j.gpb.2021.03.009.

