## Supplementary material for "Decoding human *in vitro* terminal erythropoiesis originating from umbilical cord blood mononuclear cells and pluripotent stem cells": SUPPLEMENTAL MATERIALS-20221224.docx

**Table S1 Primers for quantitative real-time PCR**

| **Target gene** | **sequence** |
| --- | --- |
| 18S-F | AACCCGTTGAACCCCATT |
| 18S-R | CCATCCAATCGGTAGTAGCG |
| HBA2-F | TCTCCTGCCGACAAGACCAA |
| HBA2-R | GCAGTGGCTTAGCTTGAAGTTG |
| HBB-F | AGGAGAAGTCTGCCGTTACTG |
| HBB-R | CCGAGCACTTTCTTGCCATGA |
| HBM-F | TCTTCACGGTGTACCCCAG |
| HBM-R | ATTAGCAGCGGAAAGTTGGCT |
| HBD-F | GACTGCTGTCAATGCCCTGT |
| HBD-R | AAAGGCACCTAGCACCTTCTT |
| HBE1-F | ATGGTGCATTTTACTGCTGAGG |
| HBE1-R | GGGAGACGACAGGTTTCCAAA |
| HBG-F | TGGATGATCTCAAGGGCAC |
| HBG-R | TCAGTGGTATCTGGAGGACA |

**Supplementary Methods**

**Single-cell gene expression quantification and cluster detection.**

Single-cell dissociations were rapidly thawed, washed and loaded into the Chromium system (10x Genomics) targeting 5,000 cells. Following the manufacturer’s instructions, barcoded sequencing libraries were generated using Chromium Single Cell 3′ v3 Reagent Kits and sequenced across 8 lanes on an Illumina NovaSeq 6000 system targeting 100,000 reads per cell. Sequencing data were aligned to the human reference genome (GRCh38) and processed using the CellRanger 4.0.0 pipeline (10x Genomics). The raw gene expression matrix from the CellRanger pipeline was imputed by the R package scImpute (version 0.0.9) (65), filtered, normalized using the R package Seurat (version 4.0.0) (66), and selected according to the following criteria: cells with >500 unique molecular identifier (UMI) counts; >200 genes, and < 25% mitochondrial gene expression in UMI counts. From the filtered cells, the gene expression matrices were normalized to the total UMI counts per cell and transformed to the natural log scale. For batch correction, we used harmony implemented in Seurat. Variable genes were selected by the function FindVariableFeatures as the top 2,500 highly variable genes expressed by more than 0.1% of cells in each sample. The R package DoubletFinder (version 2.0.3) was used to remove double cells(67), assuming a 7.5% doublet formation rate - tailor for our data. We aligned the harmony subspaces, clustered, and visualized the aligned harmony using UMAP projection. The harmony resolution was selected variably per sample origin. The major cell types were annotated according to HPCA and published reference dataset by the R package SingleR (version 1.4.1) (19) . Cell phases analysis was performed by R package scran (version 1.18.5) (31).

**DEGs identification**

DEGs were identified by performed the function ‘FindAllMarkers’ or ‘FindMarkers’ in Seurat using the ‘bimod’. The threshold of DEGs were adjusted P value is equal to 0.05 and log2 fold change is equal to 0.25.

**Pseudo-time trajectories analysis**

The R package Monocle (version 2.20.0) was used to illustrate cell state transitions during erythropoiesis(20); it applies a reversed graph embedding technique to reconstruct single-cell trajectories. We converted Seurat object to create a celldataset object using the as.CellDataSet function contained in the R package Seurat by the default settings. Monocle variable genes were used with the following cutoff criteria. The DDRTree method and the orderCells function was used to perform the construction of pseudo-time trajectories.

**Cell–cell interaction analysis**

Intercellular information is transmitted through the interaction of a cell surface receptor and through a small soluble molecule, cytokines. Using the R package CellChat (version 1.1.2) to clearly observe the interaction between each cell group (68). First, it is necessary to obtain the expression matrix and grouping information in the Seurat object, to compare the results with the “Cell-Cell Contact” database and the “Secreted Signaling” database, and then to obtain the information about the ligand and receptor interaction in pathways and core genes.

**Functional enrichment analysis**

The GO terms and gene set enrichment analysis was performed by the R package clusterProfiler (version 3.18.1) (69). The differences in pathway activities scored per cell between different cell groups were calculated with the R package GSVA (version 1.38.2) (70).

**Antibody Staining, FACS, and Flow Cytometry**

1×10^6^/test cells were prepared for flow cytometry analyses. Antibodies were incubation in 4℃ for 30min, analyze the protein expression after wash once with PBS. Data were analyzed by Flowjo (version 10, NJ, USA).

**Cell Counting Kit-8 proliferation Assay**

Seed 2000 cells in 100µL medium on every well of 96-well plate, proliferation metabolism were tested every 24 hrs. Proliferation were measure by OD450 (SpectraMax Mze, Molecular Devices, USA) after incubation cells with 10µL CCK8 (CK04-20, Dojindo, Japan) for 37℃. Data were recorded by SoftMax Pro 6.

**Real-time PCR**

Described cells were rinsed in Trizol (invitrogen, USA) for total RNA extraction, then reverse transcript to cDNA by kit (TOYOBA, Japan). To explore gene transcript level during erythropoiesis, real-time PCR were applied based on primers as table S1. Data were analyzed by GraphPad Prism (version 9.0.0, USA).

**Figure S1. The information of four samples and the annotation, distribution, and quality of cell groups**

(A-B) strategy of generation erythroid cells from PSC(A) and UCBMN(B).

(C) Morphology develop in PSC of (B).

(D) The dot plot showing the numbers and percentages of 4 samples derived from PSC and UCBMN, respectively.

(E) UMAP atlas of the PSC derived and UCBMN derived cells labeled with 4 samples.

(F) Heatmap showing the Spearman correlation coefficient between 4 samples.

(G) The density dot plot showing the Spearman correlation that calculated by all highly quality genes between PSC derived and UCBMN derived cells.

(H) Heatmap representing the cell scores of 16 cell types annotated by SingleR. PSC derived cells are colored by blue and UCBMN derived cells are colored by pink.

(I) UMAP atlas showing the cell populations in two origin derived cells annotated by SingleR.

(J) UMAP atlas showing the location of each cell type in two origins derived cells annotated by SingleR.

(K) The box plot showing the counts (top) and the features (bottom) of each cell type in Figure. 1A.

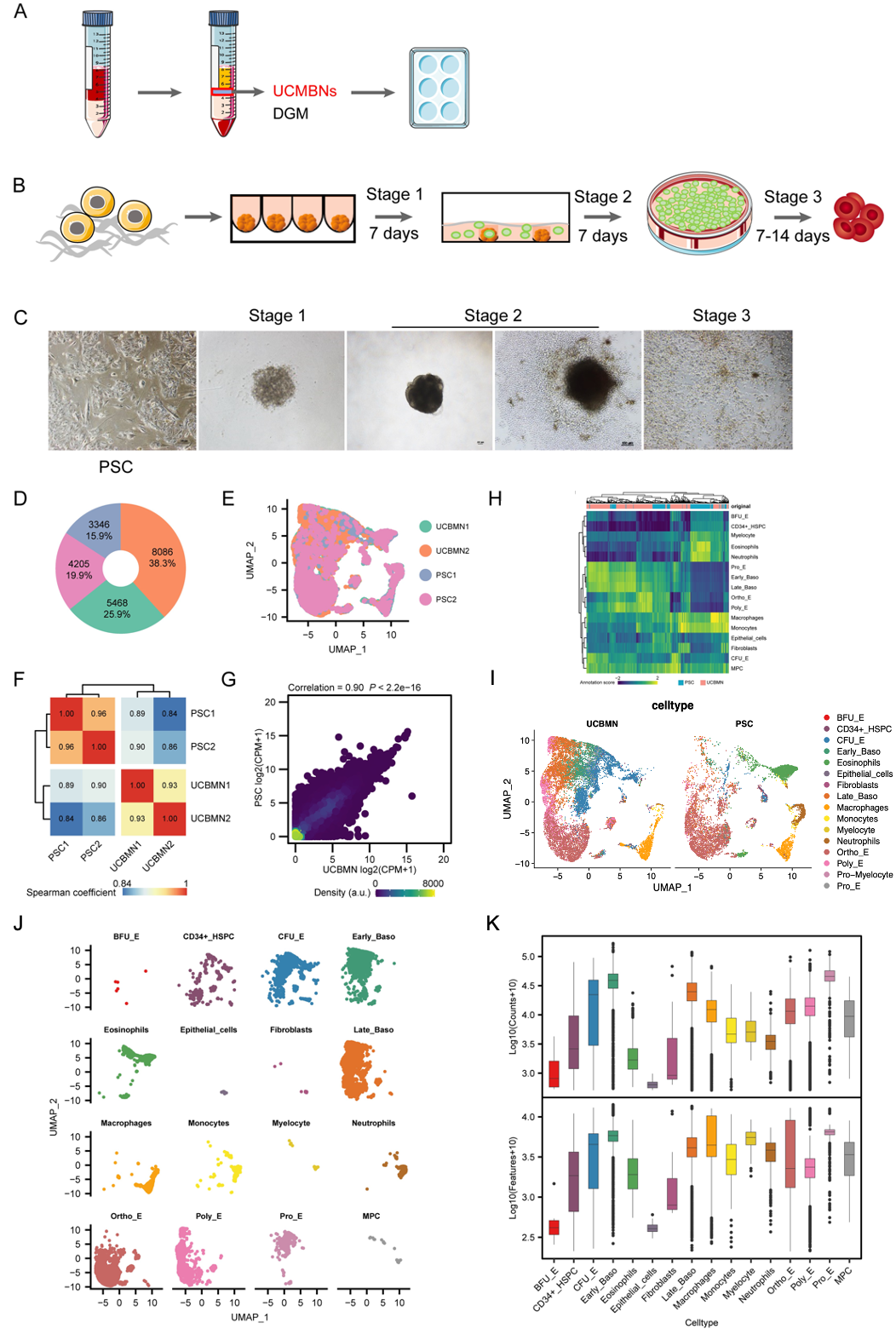

**Figure S2. The difference of early/late path in cell origins**

(A-B ) The developmental pseudo-time trajectories of progenitor and erythroid cells in different cell origins, colored by the score of pseudo-time.

(C) The pie plot showing the cell numbers of early and late paths in different cell origins.

(D) Heatmap showing the Spearman correlation coefficient between the early and late paths in different cell origins.

(E) Venn plot showing the overlap of top 20 marker genes (ordered by log2 fold change, P value < 0.05) that representing in Figure. 2B and Figure. 2C between the each early and late path in different cell origins.

(F) Venn plot showing the overlap of regulons that representing in Figure. 3A between different cell origins.

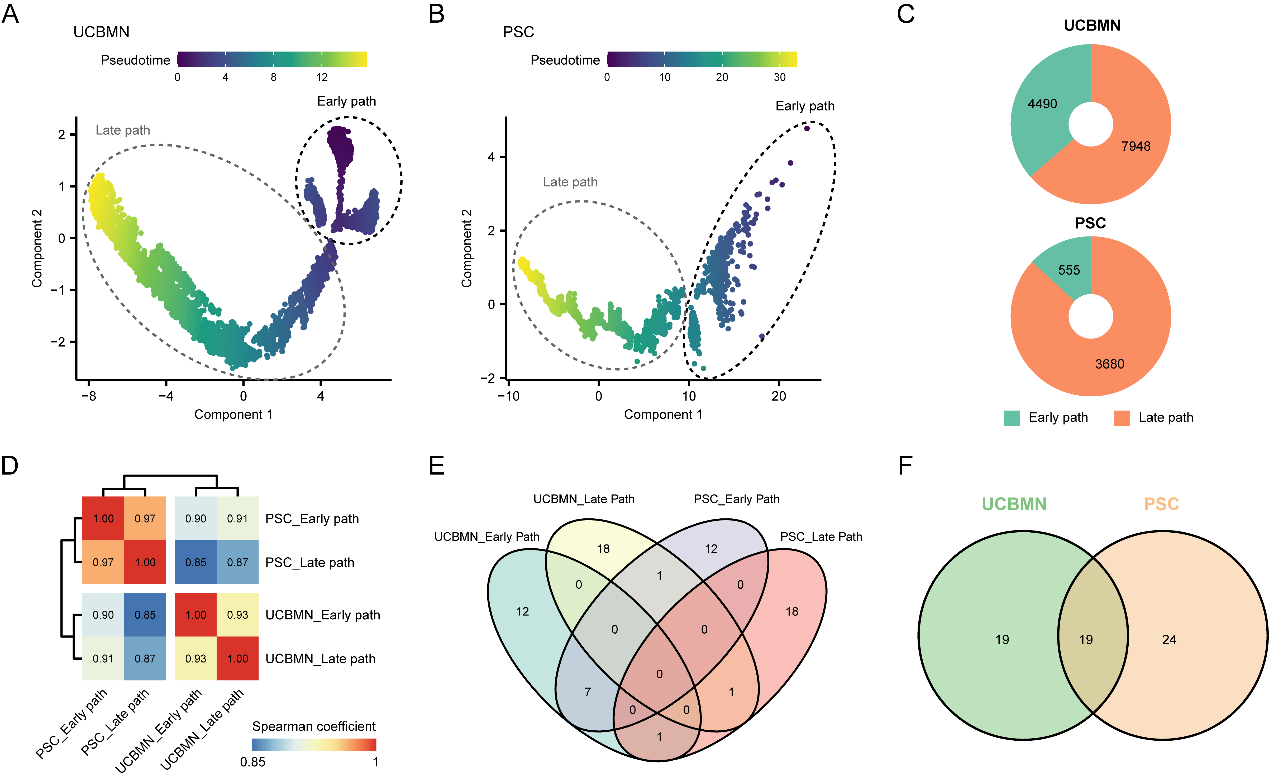

**Figure S3. The hemoglobin expression and cells construction of erythroid cells**

(A) UMAP atlas representing the distribution of adult, fetal, embryonic, and total globin.

(B)The statistical significance of each expressed globin type between differentially derived progenitor and erythroid cells in Figure 4B. ***: P value < 0.001.

(C) The pie plot representing the numbers and proportion of each cell type that showing at Figure. 4A in different cell origins.

(D) GO enrichment analysis results of genes showing in Figure. 4D.

(E) Heatmap representing the Spearman correlation coefficient of genes that showing in Figure. 4D in PSC derived (left) and UCBMN derived ortho-E cells (right), respectively.

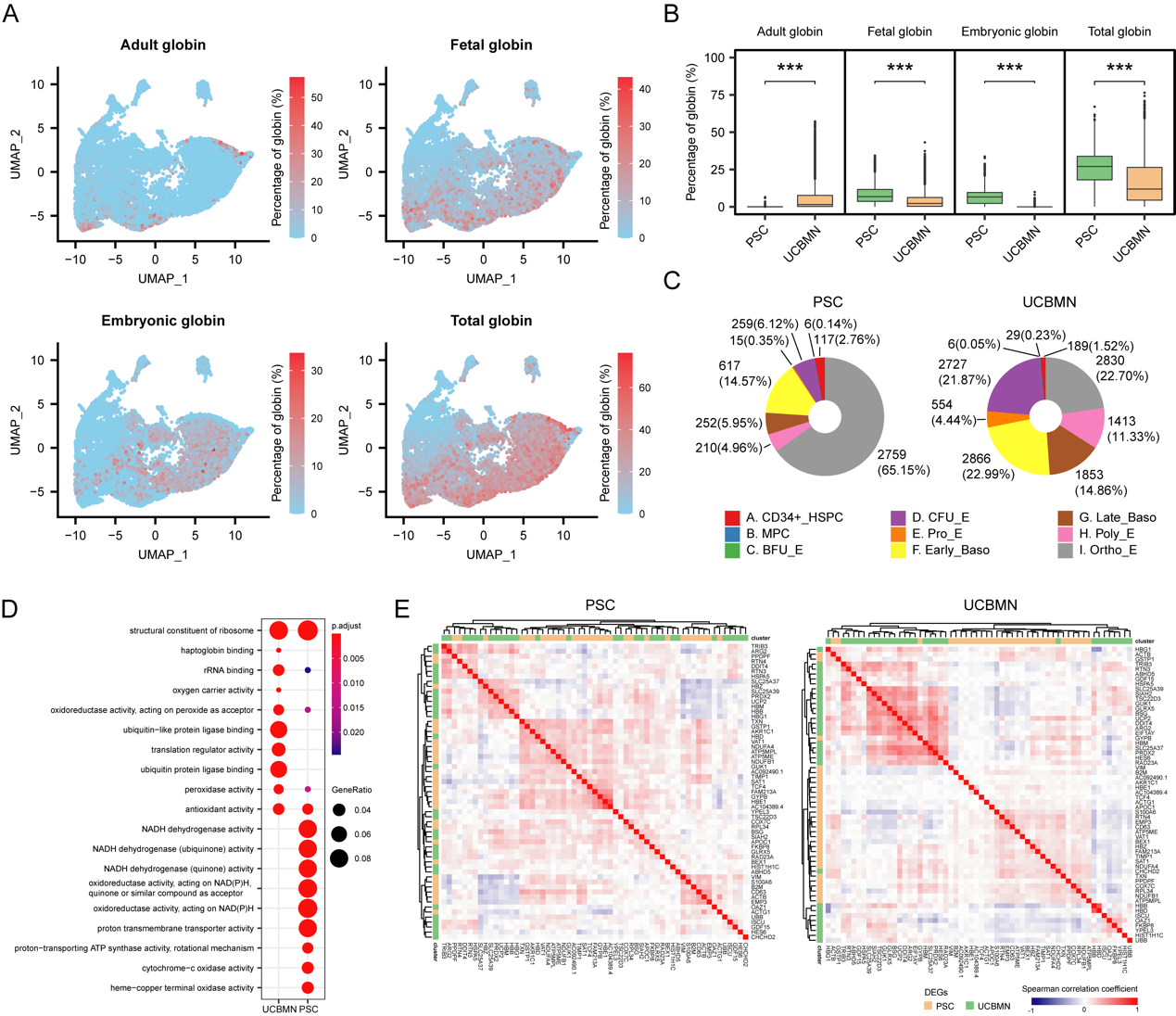

**Figure S4. The difference between early and late CFU-E cells**

(A) Violin plot showing the count number (top) and the feature number (bottom) of each progenitor and erythroid cells from different cell origins.

(B) UMAP atlas of PSC derived and UCBMN derived progenitors and erythroid cells labeled with CFU-E and other cells.

(C) Donut plot showing the cell number and percentage of CFU-E from different cell origins.

(D) K-means clustering for PSC derived (left) and UCBMN derived (right) CFU-E cells.

(E) The violin and box plot representing the count numbers of sub clustered CFU-E cells for UCBMN derived (left) and PSC derived (right) CFU-E cells.

(F) The violin and box plot representing the feature numbers of sub clustered CFU-E cells for UCBMN derived (left) and PSC derived (right) CFU-E cells.

(G) UMAP atlas representing the distribution of CFU-E cells labeled with cell origins (left) and k-means sub clusters (right).

(H) The bar plot representing the cell proportion of cell origins for k-means sub CFU-E clusters.

(I) The developmental pseudo-time trajectory of CFU-E cells labeled by cell origins. Arrow represents the direction of differentiation.

(J) Heatmap representing the pseudo-time expression of top 20 marker genes (ordered by averaged log2 fold change) that differentially expressed between k-means sub CFU-E cluster 2 and k-means sub CFU-E cluster 1.

(K) The bar plot representing the proportion of cell cycle phases for early and late CFU-E subsets.

(L) The dot plot representing the differentially expression of transcription factors between early and late CFU-E subsets. Color indicates the scaled average expression and dot size indicates the percentage of cells that expressed transcription factors in each CFU-E subset.

(M) Representative flow histogram of CD99^high^ and CD99^low^ subpopulations in CFU-E cells.

(N) Giemsa staining of FACS sorted CD99^high^ and CD99^low^ cells.

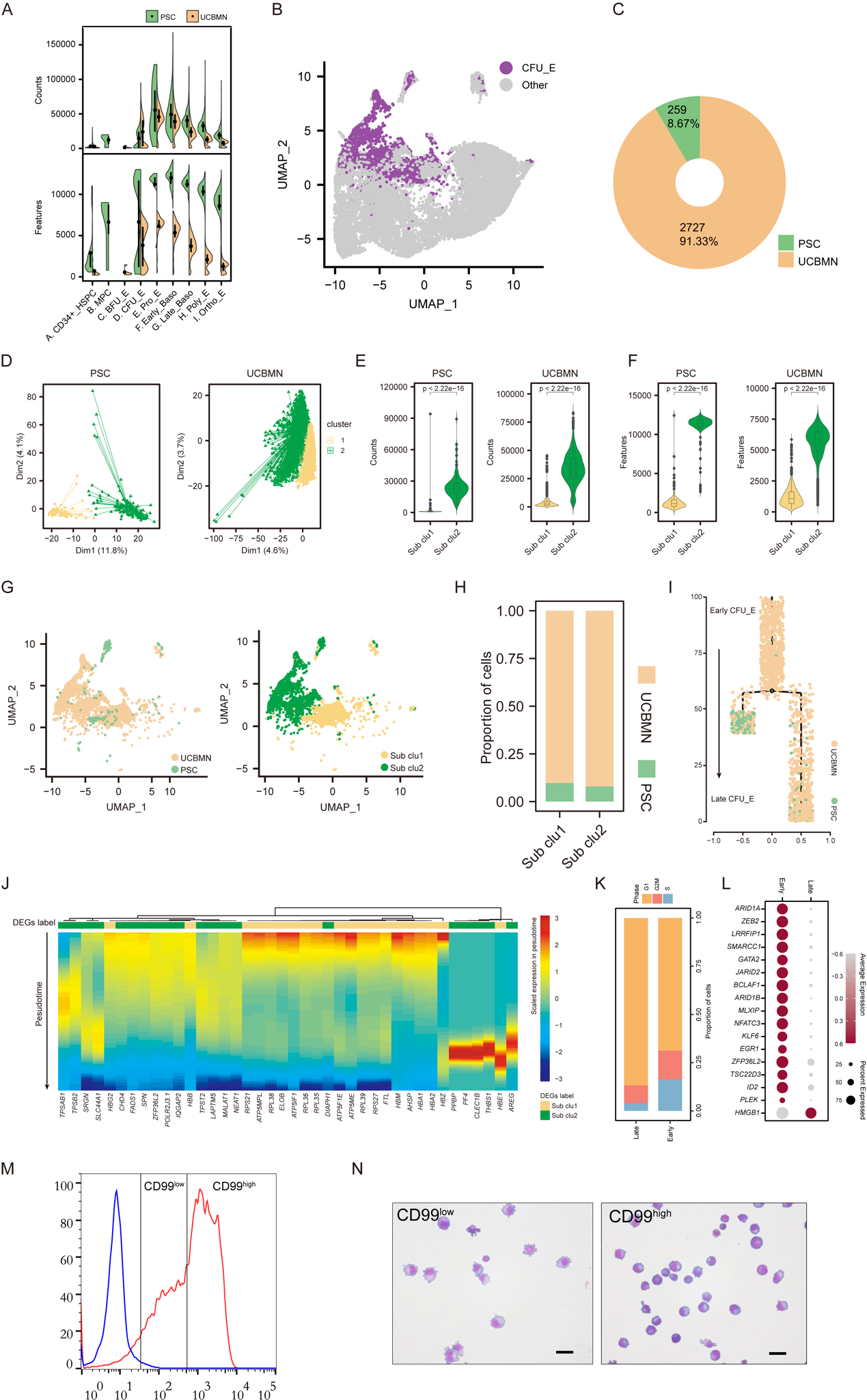

**Figure S5. CD99 marker identifies sub CFU-E cells in naïve BM CD34^+^ sample**

(A) K-means clustering for bone marrow CD34 positive (BM CD34^+^) CFU-E cells.

(B) UMAP atlas representing the distribution of BM CD34^+^ CFU-E cells labeled with k-means sub clusters.

(C) The violin and box plot representing the count numbers, the feature numbers, and the scaled CD99 expression level of sub clustered BM CD34^+^ CFU-E cells.

(D) The developmental pseudo-time trajectory of BM CD34+ CFU-E cells labeled by pseudo-time scores (top) and sub K-means clusters (bottom).

(E) The GO terms of DEGs set between highly CD99 expressed sub CFU-E cluster (top) and low CD99 expressed sub CFU-E cluster (bottom) in BM CD34+ CFU-E cells.

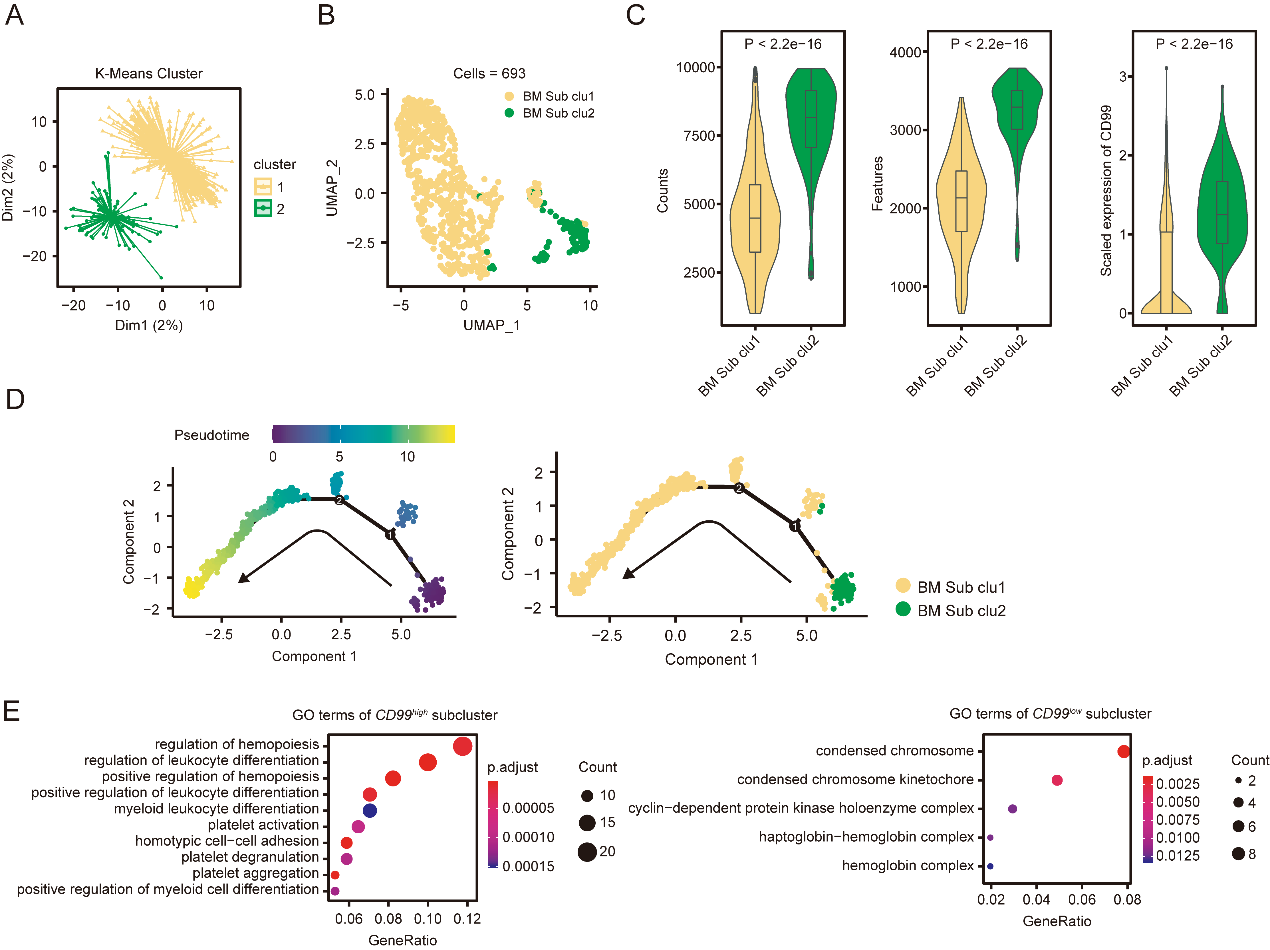

**Figure S6. Schematic model of macrophage contacting with CFU-E during the maturation of erythroid cells *in vitro***

(A) The cell-cell communication network showing the number of interactions between two cell groups at cell-cell contact pattern.

(B) The dot plot showing the significant ligand-receptor pairs from the early and the late CFU-E cells to macrophages and monocytes (left) and from macrophages and monocytes to the early and the late CFU-E cell at cell-cell contact pattern (right). Dot size represents the statistical significance of ligand-receptor pairs, and color represents the communication probability of ligand-receptor pairs.

(C) The bar plot showing the relative contribution of each ligand-receptor pair to the overall CD99 signaling pathway.

(D) CD99 expression in UCBMN-derived CFU-E were gradually decreased during in vitro erythropoiesis. The CD99 expression level in UCBMN-derived CFU-E cells were tested in early(D6), medium (D14) and late stage (D21) of erythroid production by flow cytometry analysis.

(E) The violin plot showing CD99 expression in derived erythroid cell populations.

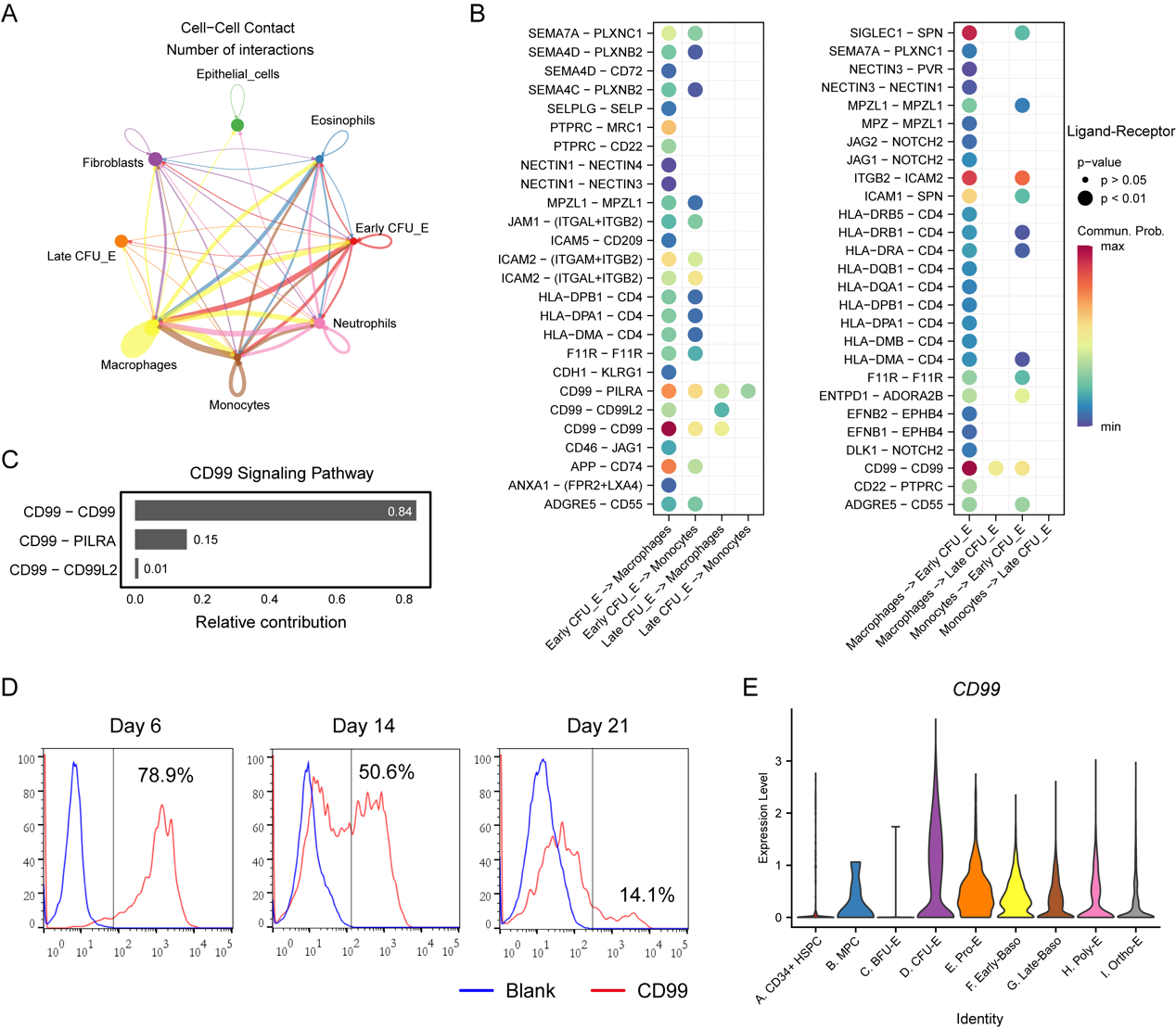
